# Distinctive Functionality of Semantic Hubs: Evidence from Semantic Dementia

**DOI:** 10.1101/2025.11.06.686766

**Authors:** Zhenjiang Cui, Di Liu, Zhiyun Dai, Xuliang Zhang, Jiahong Zeng, Qihao Guo, Zaizhu Han, Lin Huang

## Abstract

Semantic cognition relies on a distributed network with multiple candidate hub regions that bind multimodal conceptual information. While the anterior temporal lobe (ATL) is widely regarded as a core semantic hub, the roles of other candidate regions—such as the fusiform gyrus (FFG) and posterior middle temporal gyrus (pMTG)—remain debated, particularly regarding verbal and non-verbal semantic processing within the same sensory modality. This study examined 33 patients with semantic dementia and 20 healthy controls using a comprehensive behavioral battery (word and picture versions of the Pyramid and Palm Trees Test and a word–picture verification task) combined with multimodal neuroimaging, including voxel-based morphometry, diffusion-weighted imaging, and resting-state fMRI. Partial correlation and commonality analyses revealed that (1) the left ATL selectively underpins abstract verbal semantic processing, (2) the left FFG contributes to general semantic processing across verbal and non-verbal stimuli, (3) and the left pMTG plays a critical role in integrating verbal and non-verbal semantic information. White matter tracts linking the left ATL and FFG to the right ATL significantly predicted semantic integration performance, underscoring the importance of inter-hemispheric structural connectivity. Functional measures in contralateral regions, including amplitude of low-frequency fluctuation (ALFF) in the right FFG and degree centrality (DC) in the right pMTG, also predicted integration outcomes, suggesting compensatory network reorganization. These findings highlight functional specialization among semantic hubs and underscore the importance of multimodal, cross-hemispheric approaches for understanding the neural architecture of semantic representation.

## Introduction

Semantic information (i.e., the conceptual meaning of an object) refers to the abstract knowledge that allows us to recognize, categorize, and understand objects, words, and events. It plays a crucial role in everyday life, learning, and professional activities, allowing humans to understand language, recognize objects, and interact meaningfully with the world (Bechtold et al., 2023; Irish & Piguet, 2013; Liu et al., 2025; Pexman et al., 2013). How semantic information is represented and stored in the human brain remains a central question in neuroscience (Binder et al., 2009; Binder & Desai, 2011; Kumar, 2021). According to the “hub-and-spoke theory”, semantic knowledge is distributed across various modality-specific regions of the brain, commonly referred to as “spokes”. In addition, there is a cross-modal integration hub whose role is to combine information from different modalities (such as vision, audition, and language) into an abstract, amodal semantic representation. Among the numerous brain regions, the anterior temporal lobe (ATL) is widely recognized as a hub for semantic representation (Chen et al., 2020; Chiou & Lambon Ralph, 2019; Mollo et al., 2017; Muraki et al., 2025; Patterson & Lambon Ralph, 2016; Xiao et al., 2024). This region is closely associated with abstract semantic representations, and damage to it leads to impairments in semantic processing across multiple modalities (Han et al., 2013; Reilly & Peelle, 2008; Visser et al., 2010; Wang et al., 2023; Zhao et al., 2017).

However, despite the strong evidence supporting the ATL as a core hub, this view is not without controversy. Some researchers argue that semantic processing may rely on a distributed network of multiple candidate hub regions—such as the posterior middle temporal gyrus (pMTG) and the fusiform gyrus (FFG)—rather than a single amodal hub in the ATL (Chen et al., 2020; Davey et al., 2016; Forseth et al., 2018; Petrides, 2023). Evidence from neuroimaging, electrophysiology, transcranial magnetic stimulation (TMS), or lesion studies suggests that these two regions may also serve specialized roles in semantic integration and representation, thereby challenging the traditional notion of a single semantic hub (Davey et al., 2015; Ding et al., 2016; Hoffman et al., 2012; Tong et al., 2022; Tune & Asaridou, 2016; Zhang et al., 2016; Zhao et al., 2021). Moreover, while the hub-and-spoke framework emphasizes the integration of information from different modalities, it remains unclear how the brain processes different types of stimuli within the same sensory modality. Although both pictures and words are visually perceived, they engage distinct semantic processing pathways—non-verbal object semantics and verbal semantics, respectively (Bezsudnova et al., 2024; Butler et al., 2009; Federmeier & Kutas, 2001; Kahlaoui et al., 2007; Nelson & Castano, 1984; Senaha et al., 2007; Willems et al., 2008). Whether these two types of stimuli ultimately converge on a common hub or rely on partially distinct sub-hubs remains an open empirical question. Addressing this issue is critical for elucidating the neural architecture of semantic representation and may help explain why multiple candidate hubs have been identified within the temporal lobe (Clarke, 2020; Li et al., 2024; Snowden et al., 2018; Xu et al., 2017; Zhang et al., 2024).

Although existing research has provided valuable insights into the processing of verbal and non-verbal semantic information, the relationship between semantic processing of different stimulus types and the potential semantic hubs within the temporal lobe remains unclear (Butler et al., 2009; Patterson & Lambon Ralph, 2016; Uno et al., 2025; Visser et al., 2010), primarily due to the following limitations. First, direct comparisons between verbal and non-verbal semantic tasks are scarce. Many studies either focus exclusively on one stimulus type—verbal (e.g., words) or non-verbal (e.g., pictures)—or treat both as part of general semantic processing without differentiating between them. This lack of distinction makes it challenging to determine whether semantic processing for these stimuli engages a common hub or relies on distinct neural substrates (Humphreys & Lambon Ralph, 2025; Ivanova et al., 2021). Second, limitations arise from the participant groups used in previous research. Functional neuroimaging studies on healthy individuals inherently restrict causal inference, whereas lesion studies involving acute stroke patients rarely affect all three proposed semantic hubs simultaneously (Biesbroek et al., 2021; M.-M. Mesulam et al., 2019; Siddiqi et al., 2022). In particular, the ATL, which is anatomically distant from major cerebral arteries, is often spared from damage (Døli et al., 2021; Mesulam et al., 2015; Sims et al., 2016). Moreover, the widespread white matter disconnection typical of such injuries complicates efforts to isolate the specific contributions of individual gray matter regions to semantic processing (Han et al., 2013; Matchin et al., 2022; Sierpowska et al., 2019). Third, the FFG poses unique methodological challenges due to its ventral temporal location, which limits the feasibility of virtual lesion techniques such as TMS for directly probing its functional role (Brunyé et al., 2017; Rangarajan et al., 2014; Riddle et al., 2022). These limitations have prevented researchers from systematically examining and comparing, within the same participants, the selective and causally necessary contributions of multiple potential semantic hubs in the temporal lobe to verbal and non-verbal semantic processing.

Patients with semantic dementia provide a unique opportunity to address these limitations because of the nature of their progressive atrophy. In this population, atrophy typically begins in the ATL—more severely in the left hemisphere—and gradually extends to adjacent temporal regions (Czarnecki et al., 2008; Mesulam et al., 2014; Snowden et al., 2018), often in a bilateral but asymmetric pattern (Brambati et al., 2009; Woollams & Patterson, 2018). Patients with semantic dementia exhibit impairments in both verbal semantic tasks (e.g., word comprehension) and non-verbal semantic tasks (e.g., object recognition) (Bozeat et al., 2000; Corbett et al., 2009; Mesulam et al., 2013). Given the selective and progressive involvement of the temporal lobe, semantic dementia serves as an ideal lesion model for systematically examining the selective contributions of multiple candidate semantic hubs to verbal and non-verbal semantic processing (Chen et al., 2017; Chen et al., 2019; Ding et al., 2020; Lambon Ralph et al., 2017; Sundqvist et al., 2020). Moreover, previous studies primarily focused on the functional role of the ATL in this population, providing detailed and in-depth investigations, whereas relatively little attention has been paid to more posterior regions such as the pMTG and the FFG. This research bias has left our understanding of the semantic processing network incomplete (Bonner & Price, 2013; Chen et al., 2018; Davey et al., 2015; Ding et al., 2016; Liu et al., 2025; Xu et al., 2016). To better elucidate the distinct and overlapping contributions of the ATL, pMTG, and FFG to the semantic network, it is crucial to further compare their roles in verbal and non-verbal semantic processing within this patient population.

To fill above gap, in the present study, we investigated a large sample of semantic dementia patients to explore the functions of the three hubs concurrently, using a comprehensive battery of cognitive tasks: the picture version of the Pyramid and Palm Trees Test (pPPT), which evaluates non-verbal semantic processing; the word version of the PPT (wPPT), which assesses verbal semantic processing; and a word–picture verification (WPV) task, designed to examine semantic matching integration between verbal and non-verbal representations.

Multimodal neuroimaging data from patients with semantic dementia—including T1-weighted structural images, Diffusion-weighted imaging data, and resting-state fMRI—were incorporated to comprehensively investigate the relationships between brain measures of multiple semantic hubs and semantic processing performance. For each hub, voxel-based morphometry (VBM) was first applied to assess the severity of regional atrophy in the bilateral ATL, FFG, and pMTG. Partial correlation analyses were conducted to evaluate the associations between atrophy in these regions and task performance, while regressing out total intracranial volume (TIV), non-semantic control tasks (sound perception, visual perception, and number proximity matching), and lower-level language processing tasks. Lower-level language tasks were defined as basic linguistic operations that primarily involve phonological and orthographic processing rather than semantic processing (Chiou et al., 2018; Leff & Lambon Ralph, 2025; Patterson et al., 2006); in this study, these tasks included word reading and repetition. To further determine each region’s unique contribution to abstract integration of verbal and non-verbal semantic information, an additional partial correlation analysis was performed with participants’ verbal and non-verbal semantic scores included as covariates. A commonality analysis was also conducted to compare the relative involvement of gray matter regions of interest (ROIs) in supporting cross-modal semantic integration. For DTI data, white matter tracts connecting semantic hubs across hemispheres were reconstructed, and partial correlation analyses were conducted to assess the predictive effects of tract integrity on semantic task performance. Collectively, multimodal neuroimaging data provided complementary insights into the differential sensitivity of semantic processing across three levels: local structure, local functional activity (as indexed by ReHo, ALFF, and fALFF), and interregional connectivity (as indexed by degree centrality and white matter tracts). Finally, for resting-state functional data, multiple indices (ReHo, ALFF, fALFF, and degree centrality) were extracted from semantic hubs in both hemispheres, and partial correlation analyses were used to examine the predictive effects of these functional measures on semantic task performance.

## Materials and methods

### Participants

The experiment involved patients with semantic dementia and healthy controls. All participants were right-handed and native Chinese speakers. Before the study began, all participants provided written informed consent. This study was approved by the Institutional Review Board of Huashan Hospital, Fudan University, and was conducted in accordance with the Declaration of Helsinki.

#### Patients with semantic dementia

A total of 33 semantic dementia patients (15 males; age: 62.27±7.49 years; years of education: 11.73±3.01) from Huashan Hospital, Fudan University, participated in the behavioral tasks and brain imaging data collection. They met the international diagnostic criteria for semantic dementia (Gorno-Tempini et al., 2011). Among them, 16 patients showed left-lateralized atrophy, and 17 had right-lateralized atrophy. All patients had normal or corrected-to-normal hearing and vision, with no history of alcoholism, head trauma, psychiatric, or other neurological disorders. The patients’ general cognitive state was assessed using the Chinese version of the Mini-Mental State Examination (MMSE; Folstein et al., 1975), with a mean score of 21.91±4.01.

#### Healthy control subjects

A total of 20 healthy control subjects were recruited through local community advertisements. Consistent with the patient group, they were also required to have normal or corrected-to-normal hearing and vision, with no history of alcoholism, head trauma, psychiatric, or other neurological disorders. Their mean MMSE score was 28.10±1.37.

The demographic variables were well matched between the patient and healthy control groups, with no significant differences in sex (χ^2^ = 0.15, *p* = 0.70), age (*t* = 1.13, *p* = 0.27), or education level (*t* = 1.52, *p* = 0.14). Additionally, the patient group exhibited significantly lower MMSE scores compared to the control group (*t* = −8.57, *p* < 0.001).

### Behavioral performance collection

This study employed three verbal or non-verbal semantic-related neuropsychological assessments: the Picture Version of the Pyramids and Palm Trees Test (pPPT), Word Version of the Pyramids and Palm Trees Test (wPPT), and Word-Picture Verification (WPV) Test. Detailed descriptions of these assessments are as follows:

#### pPPT

This test requires subjects to determine, by pressing the left or right key, which of the two pictures presented below is semantically closer to the image displayed above. This task assesses participants’ ability in non-verbal semantic processing.

#### wPPT

This test requires participants to determine, by pressing the left or right key, which of the two words presented below is semantically closer to the word displayed above. This task assesses participants’ ability in verbal semantic processing.

#### WPV

It requires subjects to judge, via key press, whether a simultaneously presented word and picture are semantically matched. This task relies on both verbal and non-verbal semantic processing, as well as the integration of the two.

Each of these three tasks consists of 50 common and familiar items from daily life, categorized into five groups with an equal number: tools (e.g., scissors, hammer), large inoperable objects (e.g., elevator, bucket), animals (e.g., monkey, rooster), vegetables or fruits (e.g., apple, potato), and other objects (e.g., umbrella, glasses).

Additionally, we collected behavioral performance data from both the patient and control groups on several non-semantic processing tasks, including sound perception (SP), visual perception (VP) and number proximity matching (NPM), as well as lower-level language processing tasks, including reading and repetition tasks. These measures were subsequently included as covariates in the partial correlation analysis to control potential confounding effects of lower-level cognitive functions and basic language processing abilities. Below is a detailed description of these tasks.

#### SP

In each trial, participants were asked to judge whether two auditory stimuli were identical. The task included a total of 44 items. This measure was included as a covariate to control for the potential influence of reduced primary auditory perceptual ability on semantic processing performance.

#### VP

Participants were asked to judge whether two black circles presented simultaneously on the screen were of the same size by pressing a key. The task consisted of 30 items. This measure was included as a covariate to control for the potential influence of reduced primary visual perceptual ability on semantic processing performance.

#### NPM

In each trial, participants saw one number displayed at the top of the screen and two numbers at the bottom. They were asked to determine, by pressing a key, which of the two bottom numbers was closer to the top number. The task included 3 items. This measure was included as a covariate to control for the potential influence of basic cognitive abilities such as numerical comparison and attentional control on semantic task performance.

#### Reading

It required subjects to read aloud a word that appeared at the center of the screen. The task consisted of 100 words divided into five categories: tools, large inoperable objects, animals, vegetables or fruits, and other objects, with 20 items in each category. This task was included as a covariate primarily to control for the potential influence of word recognition difficulties on performance in semantic processing tasks.

#### Repetition

In this task, subjects were asked to repeat a word or sentence immediately after hearing it. The task included a total of 12 items, beginning with single words and gradually increasing in difficulty, culminating in full sentence repetition. This task was included as a covariate to control for the potential influence of phonological retrieval difficulties on performance in semantic processing tasks.

All above behavioral tasks were administered in a quiet room using the DMDX program on a PC. During testing, if a subject took longer than one minute to respond to a given item, the response was marked as incorrect, and the test proceeded to the next item. Given the substantial individual differences in reaction times across tasks, accuracy was used as the behavioral performance measure for all subjects.

### Behavioral data preprocessing

Given the considerable variability in demographic characteristics among patients such age, sex, and education, their raw scores on behavioral tasks may not accurately reflect the degree of deficit. To obtain a more precise measure of impairment, we applied the single-case-to-controls method proposed by Crawford and Garthwaite (2006). Specifically, patients’ raw behavioral scores were adjusted by accounting for the performance and demographic information of 20 healthy controls and subsequently transformed into standardized t-scores. For a detailed description of this method, please refer to previous published papers (Chen et al., 2020; Han et al., 2013; Wang et al., 2020).

### Imaging data collection

All subjects were scanned using the same 3T Siemens scanner at Huashan Hospital. High-resolution 3D T1-weighted images were acquired using the magnetization-prepared rapid gradient echo (MPRAGE) sequence along the sagittal plane. The detailed parameters were as follows: repetition time (TR) = 2300 ms, echo time (TE) = 2.98 ms, flip angle = 9°, matrix size = 240 × 256, field of view (FOV) = 240 × 256 mm^2^, number of slices = 192, slice thickness = 1 mm, and voxel size = 1 × 1 × 1 mm^3^.

The sequence for diffusion-weighted images was scanned twice in the trans verse plane with the parameters: 20 diffusion weighting directions with *b* = 1000 s/mm^2^, one additional image without diffusion weighting (i.e. *b* = 0 image), repetition time= 8500 ms, echo time = 87 ms, flip angle = 90°, matrix size = 128 × 128, field of view = 230 × 230 mm^2^, slice number = 46 slices, slice thickness = 3 mm, voxel size= 1.8 × 1.8 × 3 mm^3^.

Resting-state functional images were acquired using a gradient-echo echo-planar imaging (EPI) sequence on the transverse plane with the following parameters: repetition time (TR) = 2000 ms, echo time (TE) = 35 ms, flip angle = 90°, matrix size = 64 × 64, field of view (FOV) = 256 × 256 mm², 33 axial slices with a thickness of 4 mm, and voxel size = 4 × 4 × 4 mm³. Each scan lasted 400 s, yielding 200 volumes. During scanning, participants were instructed to keep their eyes closed, remain still, stay awake, and refrain from engaging in any structured thoughts.

### Imaging data preprocessing

#### T1 imaging data

VBM was used to quantify the gray matter volume (GMV) of participants’ brains. Preprocessing of T1-weighted structural images was performed using the CAT12 (http://www.neuro.uni-jena.de/cat/; Gaser et al., 2024) toolbox in SPM12 (http://www.fil.ion.ucl.ac.uk/spm/; Ashburner, 2012). CAT12 provides a standardized pipeline for preprocessing T1 structural images, which is primarily carried out using the ‘Segment’ module. The main steps of the analysis were as follows: First, each subject’s T1 image was aligned to the average structural image using the “inverse consistency registration” algorithm. Next, the structural images were segmented into gray matter, white matter, and cerebrospinal fluid. The gray matter portion of the image is then normalized to the MNI template using the DARTEL algorithm (Ashburner, 2007). The resulting “affine + nonlinear” components from the spatial normalization are modulated, and the final gray matter images for each participant are generated. During this process, TIV of each participant is also computed and included as a covariance in subsequent linear regression models. To improve the signal-to-noise ratio and enhance the accuracy of the analysis, a Gaussian kernel with a full width at half maximum (FWHM) of 6 mm was applied for spatial smoothing of the result images.

#### Diffusion-weighted imaging data

Diffusion-weighted imaging data were preprocessed using the PANDA toolbox (Pipeline for Analyzing Brain Diffusion Images; https://www.nitrc.org/projects/panda/; Cui et al., 2013) following these steps: (i) Brain extraction: A brain mask was generated by removing non-brain tissue from the b = 0 image using the bet command in FSL (FMRIB Software Library; https://fsl.fmrib.ox.ac.uk/fsl/docs/index.html). (ii) Eddy-current and motion correction: Distortions caused by eddy currents and simple head motion were corrected by registering all diffusion-weighted images to the *b* = 0 image with an affine transformation using FSL’s *eddy_correct* command. Gradient directions were reoriented according to the resulting transformations. (iii) Diffusion tensor estimation: Diffusion tensor models were fitted to the data, and maps of fractional anisotropy (FA), axial diffusivity (AD), mean diffusivity (MD) and radial diffusivity (RD) were computed using FSL’s *dtifit* command. (iv) Spatial normalization: Individual FA images were registered to the FMRIB58_FA template (1 mm resolution) in MNI space using FSL’s *fnirt* command. The resulting warp fields were then applied to all diffusion metric maps (FA, MD, AD, RD), resampled to 2 × 2 × 2 mm³ resolution using the *applywarp* command. Finally, these metric maps were spatially smoothed using a Gaussian kernel with a FWHM of 6 mm.

#### Resting-state fMRI data

Resting-state fMRI data were preprocessed using the Data Processing Assistant for Resting-State fMRI (DPARSF) toolbox (Yan et al., 2016). To ensure signal stabilization, the first 10 functional volumes were discarded prior to formal preprocessing. Subsequent preprocessing steps included slice timing correction, head motion realignment, spatial normalization, band-pass temporal filtering (0.01–0.1 Hz), and nuisance signal regression. Additionally, the Friston 24-parameter model for head motion, as well as signals from white matter and cerebrospinal fluid, were regressed out following realignment.

After preprocessing, we computed several functional metrics: regional homogeneity (ReHo), amplitude of low-frequency fluctuations (ALFF), fractional ALFF (fALFF), and degree centrality (DC). ReHo evaluates the local synchronization of neural activity by measuring the similarity of time series among neighboring voxels. ALFF quantifies the intensity of spontaneous low-frequency BOLD fluctuations, while fALFF represents the ratio of low-frequency power to the total frequency range, providing a more robust measure against physiological noise. ALFF and fALFF values were standardized by dividing each voxel’s value by the global average, resulting in mALFF and mfALFF maps. DC reflects the functional connectivity strength of each voxel within the whole-brain network and was computed by summing Pearson correlation coefficients between each voxel and all others exceeding a threshold of r > 0.25, forming a weighted undirected adjacency matrix. All resulting metric maps were spatially smoothed using a Gaussian kernel with a full width at half maximum (FWHM) of 6 mm.

### Definition of semantic-related ROIs

#### Grey matter anatomical subdivisions

In this study, three ROIs associated with semantic processing were defined in both the left and right cerebral hemispheres, based on established anatomical templates and methodologies reported in previous literature. The left FFG was identified directly using the Automated Anatomical Labeling (AAL) template (Rolls et al., 2020). For the left ATL and the pMTG, anatomical criteria proposed by Mesulam et al (2015, 2019). were adopted. The original ATL region encompassed the temporal pole, the anterior one-third of the superior, middle, and inferior temporal gyri, the parahippocampal gyrus, and the anterior portion of the left FFG. To avoid spatial overlap with the FFG, the intersecting area between the ATL and FFG was excluded. The left pMTG was defined as the posterior one-third of the middle temporal gyrus. Figure 1 illustrates the spatial locations of these three ROIs. To obtain homologous regions in the right hemisphere, the three left-hemisphere ROIs were mirrored across the midsagittal plane.

**Figure 1:**
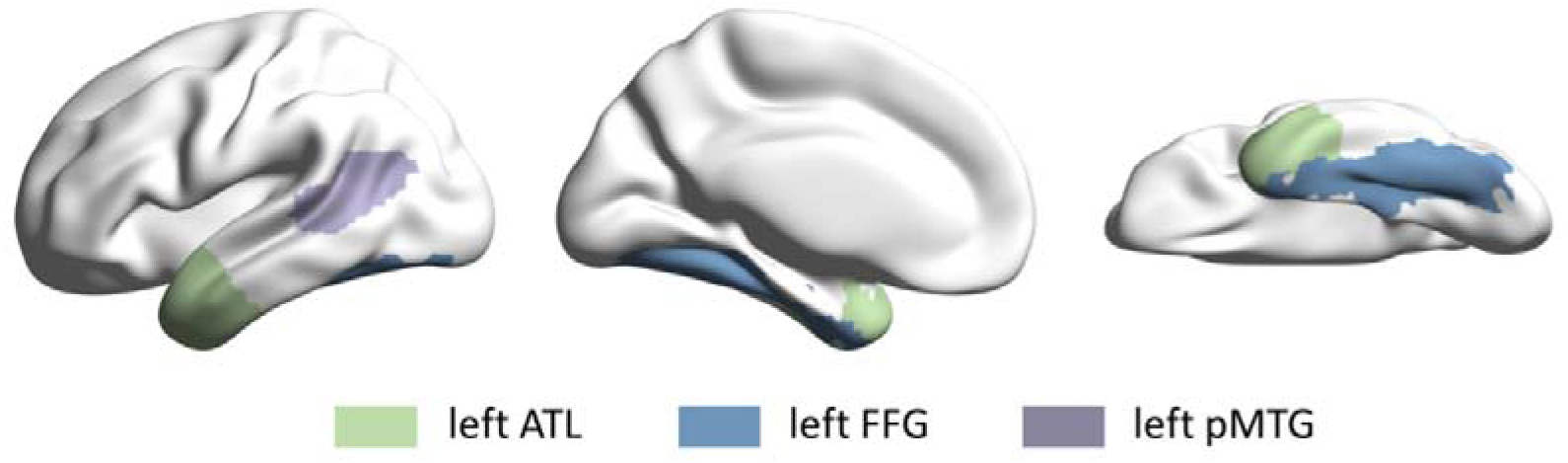
Location map of left gray matter regions of interest (ROIs). This figure shows the three left-hemisphere gray matter ROIs examined in the present study: the left anterior temporal lobe (ATL), the left fusiform gyrus (FFG), and the left posterior middle temporal gyrus (pMTG). Each region is indicated in a different color.

### White matter tractography

White matter tract masks were defined using DSI-Studio by integrating the gray matter ROIs with a standardized white matter tractography atlas constructed by Yeh and colleagues based on DWI data from 1,065 healthy participants in the Human Connectome Project (HCP). This atlas combines deterministic fiber tracking methods with manual curation by experienced neuroanatomists and clinicians to remove anatomically implausible tracts, resulting in a high-quality template of major white matter pathways (Petersen et al., 2024; Yeh, 2022). A white matter connection was considered valid only when the number of fibers between two regions exceeded one and the overlap between the generated tract mask and the gray matter ROI was less than 50% of the white matter mask voxels. The final structural connection masks were then used to extract individual-level diffusion metrics for subsequent analyses. Figure 2 illustrates the white matter connections among the six gray matter ROIs (three in each hemisphere), ultimately comprising six distinct fiber pathways linking these regions: left ATL–right ATL, left ATL–right FFG, left ATL–right pMTG, right ATL–left FFG, right ATL–left pMTG, and right ATL–right pMTG.

**Figure 2:**
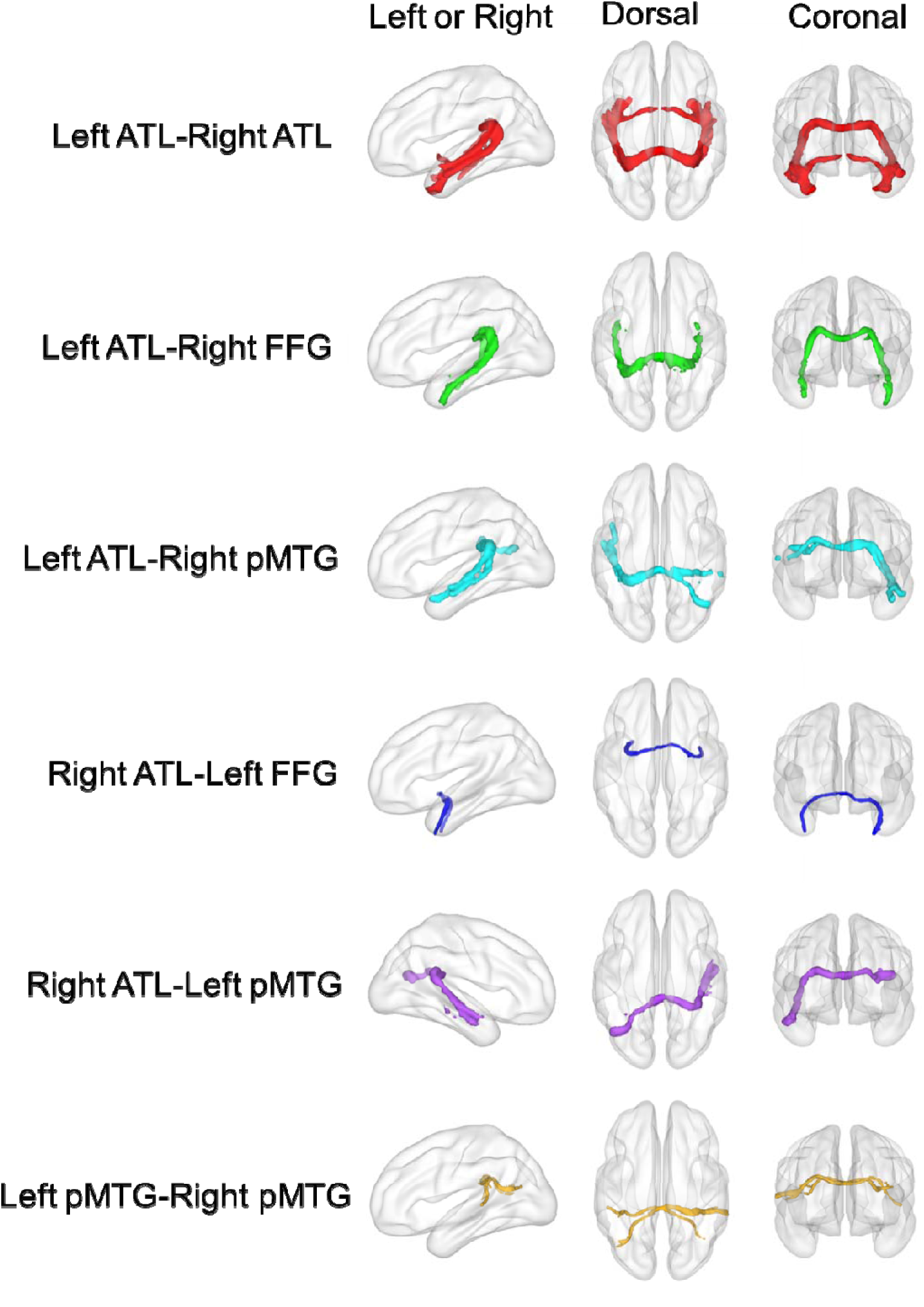
White matter tracts connecting the gray matter ROIs. Different colors indicate distinct white matter connections between brain regions, comprising a total of six fiber pathways that met the inclusion criteria.

### Statistical analysis

#### T1 imaging data

To clarify the specific contribution of grey matter structure to semantic processing, we first conducted partial correlation analyses between the gray matter volumes of three ROIs in each cerebral hemisphere and performance on three semantic tasks (pPPT, wPPT, and WPV). For each patient, mean GMV within each anatomically defined ROI was extracted by applying standard ROI masks to preprocessed T1-weighted structural images. Partial correlation analyses were then performed to assess the associations between regional GMV and behavioral performance, with TIV and performance on non-semantic control tasks (VP, SP, and NPM) included as covariates. To correct for multiple comparisons across brain regions, false discovery rate (FDR) correction was applied using the Benjamini-Hochberg procedure.

The pPPT and wPPT tasks each involve a single type of semantic information—non-verbal and verbal, respectively—whereas the WPV task requires integration of both. As such, associations between GMV and WPV performance may reflect either specific verbal or non-verbal semantic processing. To isolate the contribution of grey matter structure to abstract cross-modal semantic integration, we conducted additional partial correlation analyses between GMV and WPV performance, further controlling for pPPT and wPPT scores. In this step, none of the non-semantic control task performances were included as a covariate, as pPPT and wPPT already encompass general cognitive processing. We believed that controlling for these scores was sufficient to account for lower-level perceptual contributions.

Given the characteristic atrophy pattern in semantic dementia—where the ATL is typically the earliest and most severely affected region, with degeneration spreading to adjacent areas—correlations observed in regions such as the pMTG and FFG may be confounded by co-occurring ATL atrophy. To assess the unique contributions of these regions beyond the influence of ATL, we conducted further partial correlation analyses including the GMV of ATL as an additional covariate.

To complement these analyses, we further performed a commonality analysis to quantify the extent to which performance on the pPPT and wPPT tasks explained variance in WPV performance. The explained variance reflects modality-specific semantic contributions, while the unexplained variance is presumed to reflect abstract integrative processing across verbal and non-verbal domains. Theoretically, such integration is also supported by the semantic hub. Therefore, we further examined the unique contributions of the ATL, pMTG, and FFG to this integrative component, aiming to clarify and compare the roles of these regions in verbal and non-verbal semantic integration.

#### Diffusion-weighted imaging data

For the DWI data, we conducted partial correlation analyses within the semantic dementia group, following procedures similar to those applied to the T1 imaging data. These steps included examining correlations between white matter metrics and performance on three semantic tasks, assessing the relationship between white matter metrics and WPV task performance after regressing out TIV and scores on the pPPT and wPPT tasks, and further controlling for potentially confounding gray matter measures. Mean values of diffusion metrics (FA, AD, MD, and RD) within white matter tract masks were used as indicators of white matter connectivity integrity. The aim was to determine the extent to which the integrity of these interregional pathways predicts performance on semantic processing tasks.

#### Resting-state fMRI data

For the resting-state fMRI data, we computed regional functional indices—including ReHo, ALFF, fALFF, and DC—within three predefined gray matter ROIs in both the left and right hemispheres. Partial correlation analyses were conducted within the patient group to examine the predictive effects of these indices on performance across three semantic tasks (wPPT, pPPT, and WPV).

All brain imaging indices were extracted using MATLAB R2022a (https://www.mathworks.com/), and statistical analyses were performed using JASP version 0.95.3.0 (https://jasp-stats.org/). To correct for multiple comparisons across brain regions, the statistical results were adjusted using the Benjamini–Hochberg false discovery rate (FDR) correction method, with a significance threshold set at FDR-corrected *q* < 0.05.

## Results

### Behavioral profiles of the participants

Table 1 presents the raw scores and adjusted *t*-scores from three neuropsychological assessments for 20 healthy controls and 33 patients with semantic dementia. The adjusted *t*-score results revealed significantly worse behavioral performance across all three semantic processing tasks in patients compared to demographically matched controls. Specifically, patients showed significantly impaired performance compared to healthy controls on all three semantic tasks, including the pPPT task (mean *t*-score = −4.01, SD = 1.87), the wPPT task (mean *t*-score = −11.02, SD = 7.12), and the WPV task (mean *t*-score = −9.07, SD = 6.52). In lower-level language tasks, the patient group exhibited a significant decline in reading task performance (mean *t*-score = −9.88, SD = 10.80), whereas their performance on the repetition task remained intact (mean *t*-score = −0.96, SD = 1.69). Moreover, in non-semantic control tasks, there were no significant behavioral differences between patients and healthy controls (SP: mean *t*-score = −0.95, SD = 1.12; VP: mean *t*-score = 0.13, SD = 0.84; NPM: mean *t*-score = −0.21, SD = 1.40). These findings are generally consistent with the characteristic profile of semantic dementia, in which semantic processing is most severely affected; however, the significant impairment in reading performance suggests that lower-level language processes may also be compromised.

**Table 1.** Behavioral performance of healthy control subjects and patients.

| | Raw Score: Mean ± SD | | Corrected *t*-score ± SD |
| --- | --- | --- | --- |
|  | Healthy Controls | Semantic Dementia | of patients |
| **Semantic Tasks** |  |  |  |
| wPPT | 0.95 ± 0.03 | 0.75 ± 0.12 | -11.02 ± 7.12 |
| pPPT | 0.95 ± 0.04 | 0.75 ± 0.09 | -4.01 ± 1.87 |
| WPV | 0.96 ± 0.02 | 0.71 ± 0.18 | -9.07 ± 6.52 |
| **Lower-level** |  |  |  |
| **Language Tasks** |  |  |  |
| Reading | 0.98 ± 0.02 | 0.79 ± 0.20 | -9.88 ± 10.80 |
| Repetition | 0.98 ± 0.03 | 0.94 ± 0.07 | -0.96 ± 1.69 |
| **Non-semantic** |  |  |  |
| **Control Tasks** |  |  |  |
| SP | 0.88 ± 0.09 | 0.76 ± 0.13 | -0.95 ± 1.12 |
| VP | 0.91 ± 0.05 | 0.92 ± 0.06 | 0.13 ± 0.84 |
| NPM | 0.93 ± 0.13 | 0.87 ± 0.22 | -0.21 ± 1.40 |

Within-patient Pearson correlation analyses revealed the following relationships between semantic-related task performances: pPPT-wPPT: *r* = 0.48, *p* < 0.01; pPPT-WPV: *r* = 0.56, *p* < 0.001; wPPT-WPV: *r* = 0.82, *p* < 0.001. The results support a strong correlation between the two tasks involving word stimuli (wPPT and WPV; *r* > 0.8), indicating substantial overlap in the verbal semantic processes involved. A moderate correlation (0.4 < *r* < 0.6) was observed between the task involving only picture stimuli (pPPT) and the task involving word stimuli (wPPT and WPV). Their overlap may reflect a shared semantic system supporting both verbal and non-verbal semantic processing, while the remaining differences may be attributed to stimulus-specific or integration-related processing components.

In addition, further correlation analyses revealed that reading performance in the patient group was strongly associated with the wPPT and WPV tasks (wPPT: *r* = 0.78, *p* < 0.001; WPV: *r* = 0.67, *p* < 0.001) and showed a near-moderate correlation with the pPPT task (*r* = 0.39, *p* < 0.05). Similarly, repetition task performance showed weak-to-moderate correlations with the wPPT and WPV tasks (wPPT: *r* = 0.38, *p* < 0.05; WPV: *r* = 0.37, *p* < 0.05), but not with pPPT (*r* = 0.17, *p* = 0.36). No significant correlations were observed between non-semantic control task performance and any semantic-related tasks (|*r*s| < 0.30, *p*s > 0.10). These findings highlight the need to control for phonological and orthographic deficits when investigating abstract semantic processing, to ensure that observed effects are specific to semantic mechanisms.

### Whole-brain atrophy in patients with semantic dementia

Figure 3 displays whole-brain atrophy maps comparing 33 patients with semantic dementia to 20 healthy controls at the group level (FDR-corrected *q* < 0.05). The results reveal that patients with semantic dementia exhibit the most severe atrophy in bilateral ATL regions, extending anteriorly to the insula and frontal lobe, and posteriorly to the ventral temporal lobe, pMTG, inferior temporal gyrus (ITG) and FFG. The atrophied regions observed in the current study showed anatomical overlap with all three predefined ROIs in both the left and right hemispheres.

**Figure 3:**
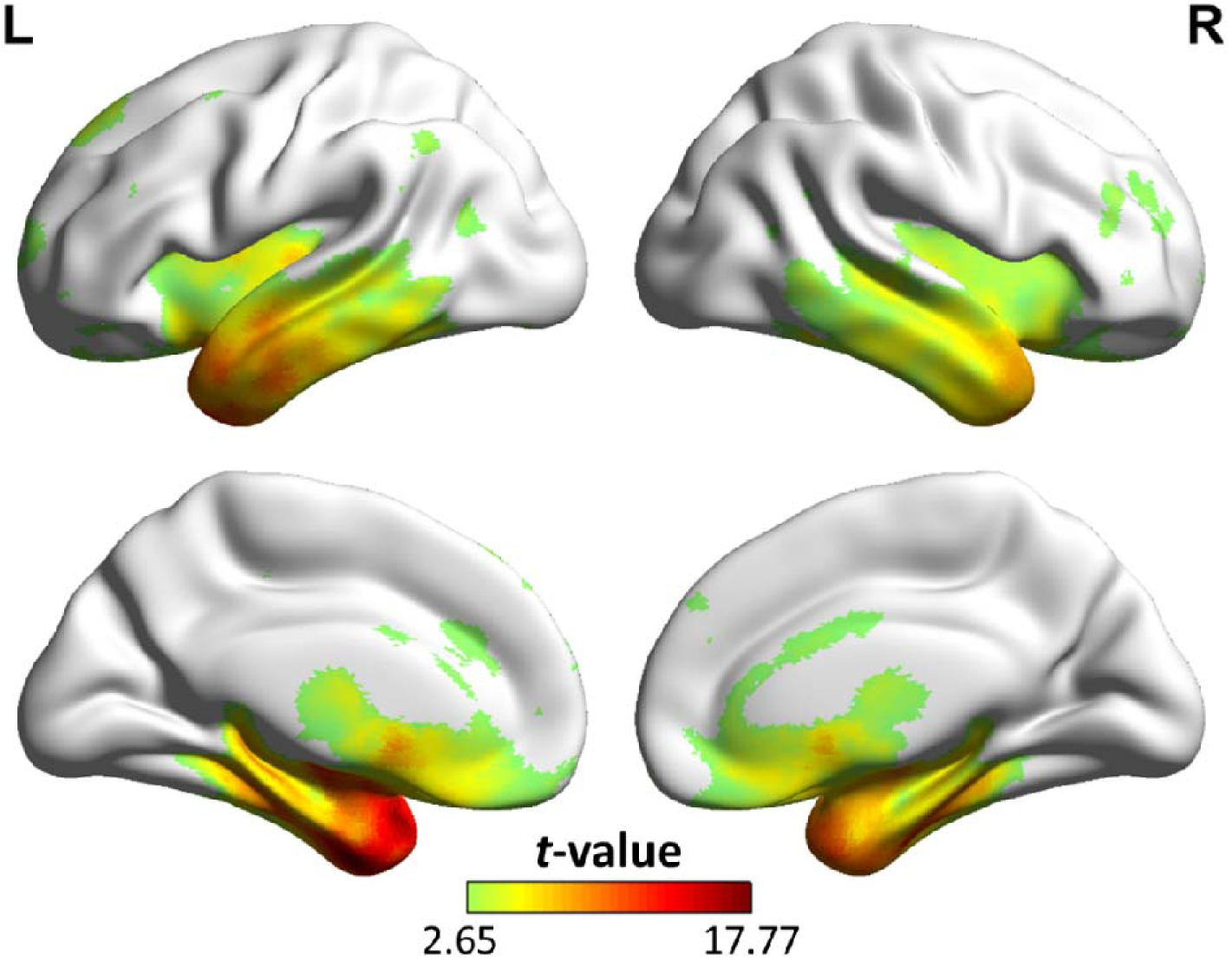
Atrophy map of the semantic dementia patients. The figure shows the areas with significant differences in GMV between the 33 semantic dementia patients and 20 healthy control subjects (FDR-corrected *q* < 0.05).

### GMV–behavior mapping analysis based on template-defined ROIs

#### ROI-based GMV–behavioral performance partial correlation analysis in the left hemisphere

After regressing out TIV, performance on non-semantic control tasks (SP, VP and NPM), as well as lower-level language processing tasks (reading and repetition), partial correlation analyses conducted within patient group revealed significant associations between behavioral performance and GMV in specific regions of left hemisphere (see Figure 4).

**Figure 4:**
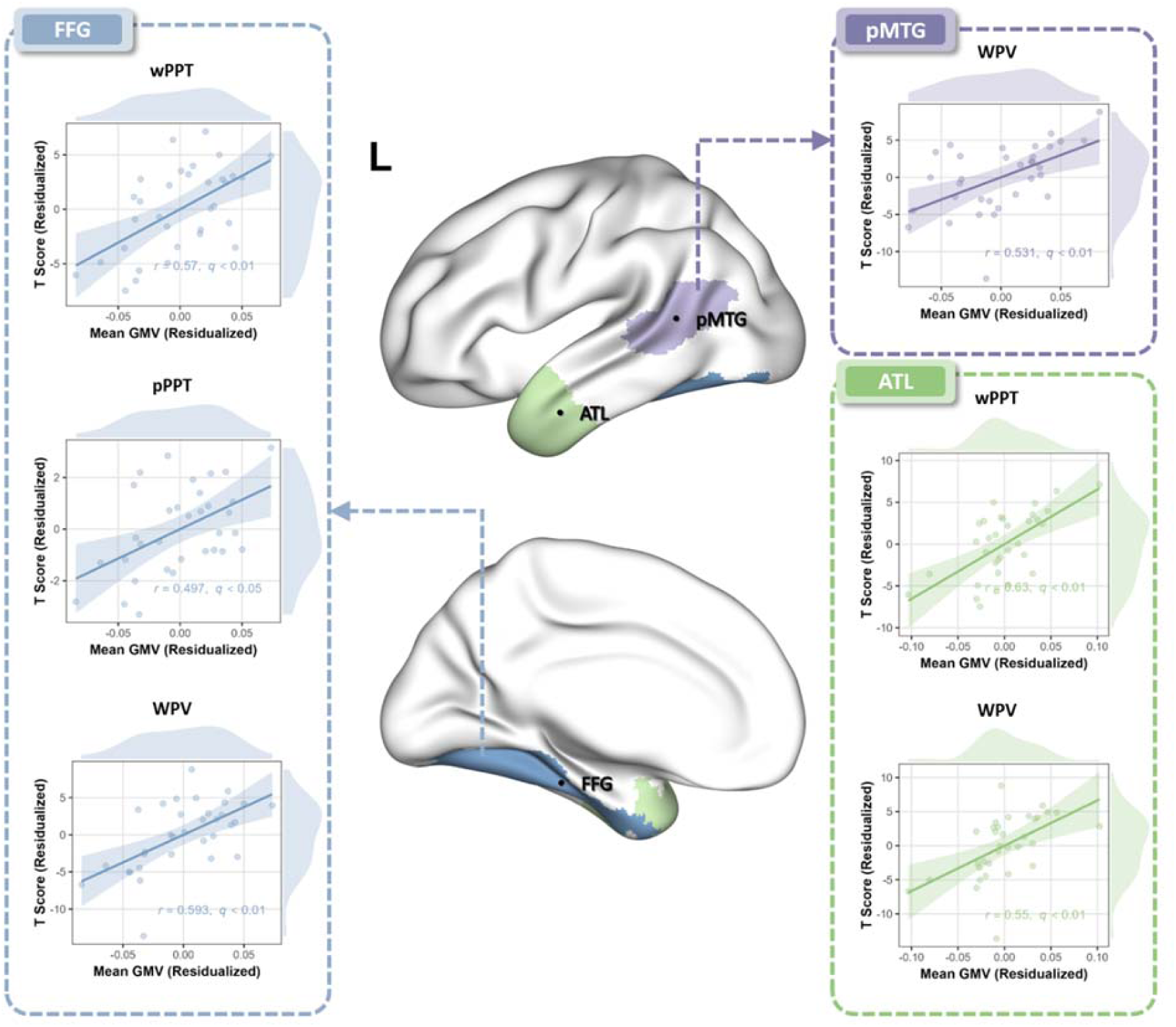
Significant correlation results between GMV in left ROIs and performance on semantic processing tasks (WPV, pPPT, and wPPT) within the semantic dementia patient group after regressing out TIV, performance on non-semantic control tasks, as well as lower-level language processing tasks. FDR-corrected *q*-values are reported.

For the wPPT task, significant correlations were found with GMV in the left ATL and FFG (*r*s = 0.63 and 0.57, FDR-corrected *q*s < 0.01). A marginally significant correlation was also found in the left pMTG (*r* = 0.407, FDR-corrected *q* = 0.035). For the pPPT task, a significant correlation was observed with GMV in the left FFG (*r* = 0.497, FDR-corrected *q* < 0.05), whereas no significant correlation was observed in the left ATL (*r* = 0.268, FDR-corrected *q* = 0.176) and pMTG (*r* = 0.312, FDR-corrected *q* = 0.170). For the WPV task, significant correlations were identified in all three regions: left ATL (*r* = 0.550, FDR-corrected *q* < 0.01), FFG (*r* = 0.593, FDR-corrected *q* < 0.01), and pMTG (*r* = 0.531, FDR-corrected *q* < 0.01).

#### ROI-based GMV–behavioral performance partial correlation analysis in the right hemisphere

After regressing out TIV, performance on non-semantic control tasks (SP, VP and NPM), as well as lower-level language processing tasks (reading and repetition), no significant correlations were observed between behavioral performance and GMV in the right ATL (wPPT: *r* = −0.051, FDR-corrected *q* = 0.833; pPPT: *r* = 0.316, FDR-corrected *q* = 0.164; WPV: *r* = 0.304, FDR-corrected *q* = 0.123), pMTG (wPPT: *r* = 0.043, FDR-corrected *q* = 0.833; pPPT: *r* = 0.258, FDR-corrected *q* = 0.193; WPV: *r* = 0.394, FDR-corrected *q* = 0.063), or FFG (wPPT: *r* = 0.106, FDR-corrected *q* = 0.833; pPPT: *r* = 0.372, FDR-corrected *q* = 0.164; WPV: *r* = 0.424, FDR-corrected *q* = 0.063) (See figure 5).

**Figure 5:**
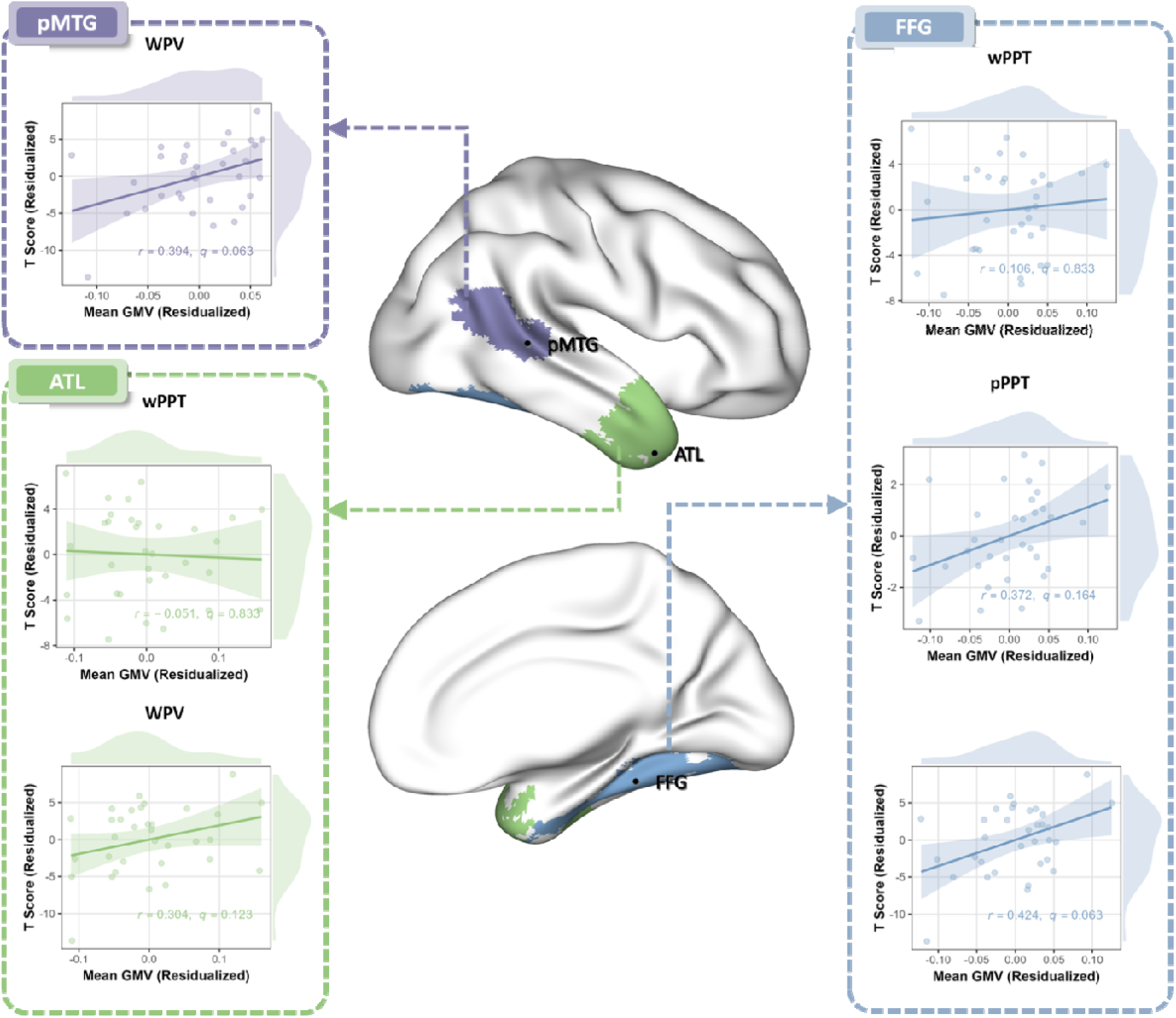
Significant correlation results between GMV in left ROIs and performance on semantic processing tasks (WPV, pPPT, and wPPT) within the semantic dementia patient group after regressing out TIV, performance on non-semantic control tasks, as well as lower-level language processing tasks. FDR-corrected *q*-values are reported.

These results indicate that semantic deficits in the patient group are primarily associated with gray matter atrophy in left hemisphere regions. Accordingly, subsequent analyses focused on examining the relationships between left hemisphere ROIs and performance on semantic tasks.

#### ROI-based GMV–behavior associations in WPV task after controlling for verbal and non-verbal semantic performance

The significant associations observed across all three left ROIs in the WPV task can also be attributed to the fact that this task engages verbal semantic processing, non-verbal semantic processing, and the integration of both. To isolate the component specifically related to cross-modal integration, verbal (wPPT) and non-verbal (pPPT) semantic performance scores were further included as covariates in a follow-up partial correlation analysis conducted within the patient group. Even after regressing out these factors, significant positive correlations remained between WPV performance and GMV in the left FFG (*r* = 0.439, FDR-corrected *q* < 0.05) and pMTG (*r* = 0.479, FDR-corrected *q* < 0.05), highlighting the critical role of these regions in supporting verbal and non-verbal semantic integration. In contrast, the correlation between the GMV of left ATL and WPV performance was marginally significant (*r* = 0.312, *FDR*-corrected *q* = 0.093). Furthermore, beyond controlling for TIV and performance on the wPPT and pPPT tasks, we additionally regressed out the GMV of their corresponding right-hemisphere (c-rGMV) regions when examining the associations between left-hemisphere ROIs and WPV performance. The results remained stable (left pMTG: *r* = 0.456, *FDR*-corrected *q* < 0.05; left FFG: *r* = 0.473, *FDR*-corrected *q* < 0.05; left ATL: *r* = 0.332, *FDR*-corrected *q* = 0.078), indicating that the observed left-hemisphere effects were not driven by shared variance with their right-hemisphere homologues (see Table 2).

**Table 2.** ROI-Based GMV–Behavior Associations in WPV Task.

|  | Regressors | *r* | FDR-corrected *q* |
| --- | --- | --- | --- |
| **Left Hemisphere** |  |  |  |
| ATL |  | 0.312 | 0.093 |
| FFG | TIV, pPPT, wPPT | 0.439 | **< 0.05\*** |
| pMTG |  | 0.479 | **< 0.05\*** |
| ATL | TIV, pPPT, wPPT | 0.332 | 0.078 |
| FFG |  | 0.473 | **< 0.05\*** |
| pMTG | c-rGMV | 0.456 | **< 0.05\*** |
| **Right Hemisphere** |  |  |  |
| ATL |  | 0.207 | 0.281 |
| FFG | TIV, pPPT, wPPT | 0.312 | 0.212 |
| pMTG |  | 0.337 | 0.207 |
| ATL | TIV, pPPT, wPPT, | 0.207 | 0.281 |
| FFG |  | 0.334 | 0.176 |
| pMTG | c-IGMV | 0.297 | 0.176 |

In addition, after regressing out the GMV of the left ATL, TIV and performance on the wPPT and pPPT tasks, the correlation between GMV in the left pMTG and performance on the WPV task remained statistically significant without FDR correction (*r* = 0.409, *p* < 0.05). In contrast, the correlation between GMV in the left FFG and WPV task performance was marginally significant (*r* = 0.339, *p* = 0.072). Furthermore, after regressing out the GMV of the left FFG, the GMV of the left pMTG showed a marginally significant association with WPV task performance within the patient group (*r* = 0.352, *p* = 0.066). However, after controlling for the GMV of the left pMTG, the GMV of the left FFG was no longer significantly associated with WPV task performance within the patient group (*r* = 0.263, *p* = 0.176).

In the right hemisphere, however, the marginally significant association between GMV in the right pMTG and FFG and performance on the WPV task disappeared after controlling for TIV, as well as performance on the pPPT and wPPT tasks (right pMTG: *r* = 0.337, *FDR*-corrected *q* = 0.207; right FFG: *r* = 0.312, *FDR*-corrected *q* = 0.212), and the correlation between the right ATL and WPV task performance remained non-significant (*r* = 0.207, *FDR*-corrected *q* = 0.281). When the GMV of the corresponding left-hemisphere (c-lGMV) regions was further regressed out, the correlation between the right pMTG (*r* = 0.297) and WPV performance slightly decreased, whereas the correlation for the right FFG (*r* = 0.334) slightly increased; however, neither correlation reached statistical significance (FDR-corrected *q*s = 0.176) (see Table 2).

The results of this part support the view that the contributions of left-hemisphere language hub regions to verbal and non-verbal semantic integration processes are largely independent of the influence of right-hemisphere hub regions, with the left pMTG and FFG taking a stronger effect on both types of integration.

#### ROI-based GMV-behavioral performance commonality analysis

Further commonality analyses within the patient group revealed the extent to which performance on the pPPT and wPPT tasks explained variance in the WPV task. Together, these two tasks accounted for approximately 70.7% of the variance in WPV performance. Specifically, the wPPT alone uniquely explained 39.0% of the variance, while the pPPT alone accounted for only 3.6%. The shared variance between the two tasks was 28.1%. In addition, the TIV contributed negligibly to the explanation of variance (<0.01%) in WPV task performance, suggesting that individual differences in brain size had minimal impact on the observed behavioral effects.

Considering that individual differences in TIV may influence the GMV of ROIs, we first constructed separate regression models with TIV as the predictor of GMV in the left ATL, FFG, and pMTG. The residuals from these models, representing GMV values adjusted for TIV, were subsequently used in all further analyses. The results showed that, after regressing out TIV, wPPT and pPPT performances, the residual GMVs of the left ATL, FFG, and pMTG further explained an additional 24.30% of the variance in WPV performance. Among these, the left pMTG accounted for the largest proportion of variance, explaining nearly half of the variance attributed to all three semantic hubs combined. In contrast, the left FFG and left ATL contributed only 1.6% and 0.1%, respectively, indicating that GMV in the left pMTG was the strongest predictor of behavioral performance (see Figure 6).

**Figure 6:**
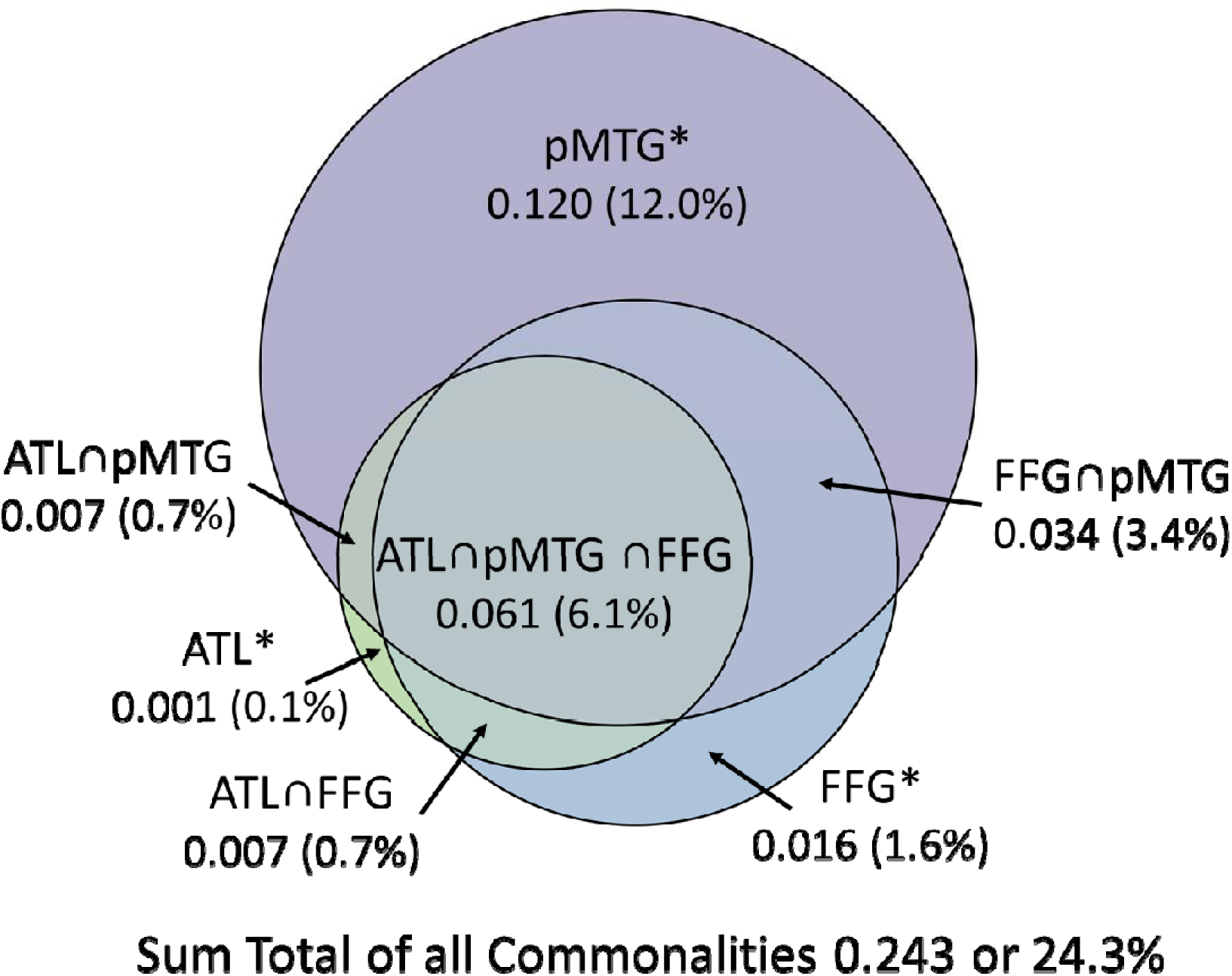
Componential analysis illustrating the unique and shared contributions of GMV in the left pMTG, left ATL, and left FFG to WPV task performance. Each circle represents one semantic hub, with overlapping areas indicating shared explained variance. Asterisks (*) denote the variance uniquely explained by each region.

These findings suggest that the left pMTG play a particularly critical role in semantic integration during the WPV task, more so than the left FFG and ATL.

### White matter connectivity–behavior mapping analysis

#### Partial correlation analysis between white matter integrity and semantic processing task performance

In the partial correlation analysis between white matter integrity and behavioral performance, results indicated that, within the patient group, FA did not significantly predict any behavioral measures (FDR-corrected *q*s > 0.10). In contrast, AD, MD, and RD consistently exhibited predictive effects on behavioral performance. Accordingly, the statistical results primarily focus on the significant predictive effects of AD, MD, and RD on behavioral performance. Furthermore, in the subsequent analysis that additionally regressed out GMV, we only performed regression on those white matter tracts that showed significant associations with behavioral performance. For these analyses, we reported the raw *p* values rather than FDR-corrected *q* values.

Within the patient group, after regressing out TIV, performance on non-semantic control tasks (SP, VP, and NPM), and lower-level language processing tasks (reading and repetition), among the six white matter tracts examined, we found that only the tract connecting the left FFG and the right ATL showed significant negative correlations between its integrity (as indexed by AD, MD, and RD) and performance on the pPPT task (AD: *r* = −0.562, FDR-corrected *q* < 0.05; MD: *r* = −0.543, FDR-corrected *q* < 0.05; RD: *r* = −0.531, FDR-corrected *q* < 0.05). Additionally, the white matter connection between the right ATL and the left FFG showed strong predictive effects on WPV performance across multiple diffusion metrics. Specifically, AD exhibited a significant negative correlation with WPV scores (*r* = −0.639, FDR-corrected *q* < 0.01). For MD and RD, significant associations were also observed (MD: *r* = −0.606, FDR-corrected *q* < 0.05; RD: *r* = −0.587, FDR-corrected *q* < 0.05), see Figure 7. Given that GMV of the left FFG was significantly correlated with performance on the pPPT and WPV tasks, we included GMV of the FFG as an additional covariate in our analyses. Even after controlling for this gray matter effect, the integrity of the white matter tract connecting the left FFG and the right ATL remained significantly and negatively associated with performance on both the pPPT (AD: *r* = −0.441, *p* < 0.05; MD: *r* = −0.438, *p* < 0.05; RD: *r* = −0.434, *p* < 0.05) and WPV (AD: *r* = −0.523, *p* < 0.01; MD: *r* = −0.508, *p* < 0.01; RD: *r* = −0.498, *p* = 0.01) tasks. These findings suggest that the integrity of this white matter pathway contributes independently to semantic processing deficits, beyond the effects of gray matter atrophy.

**Figure 7:**
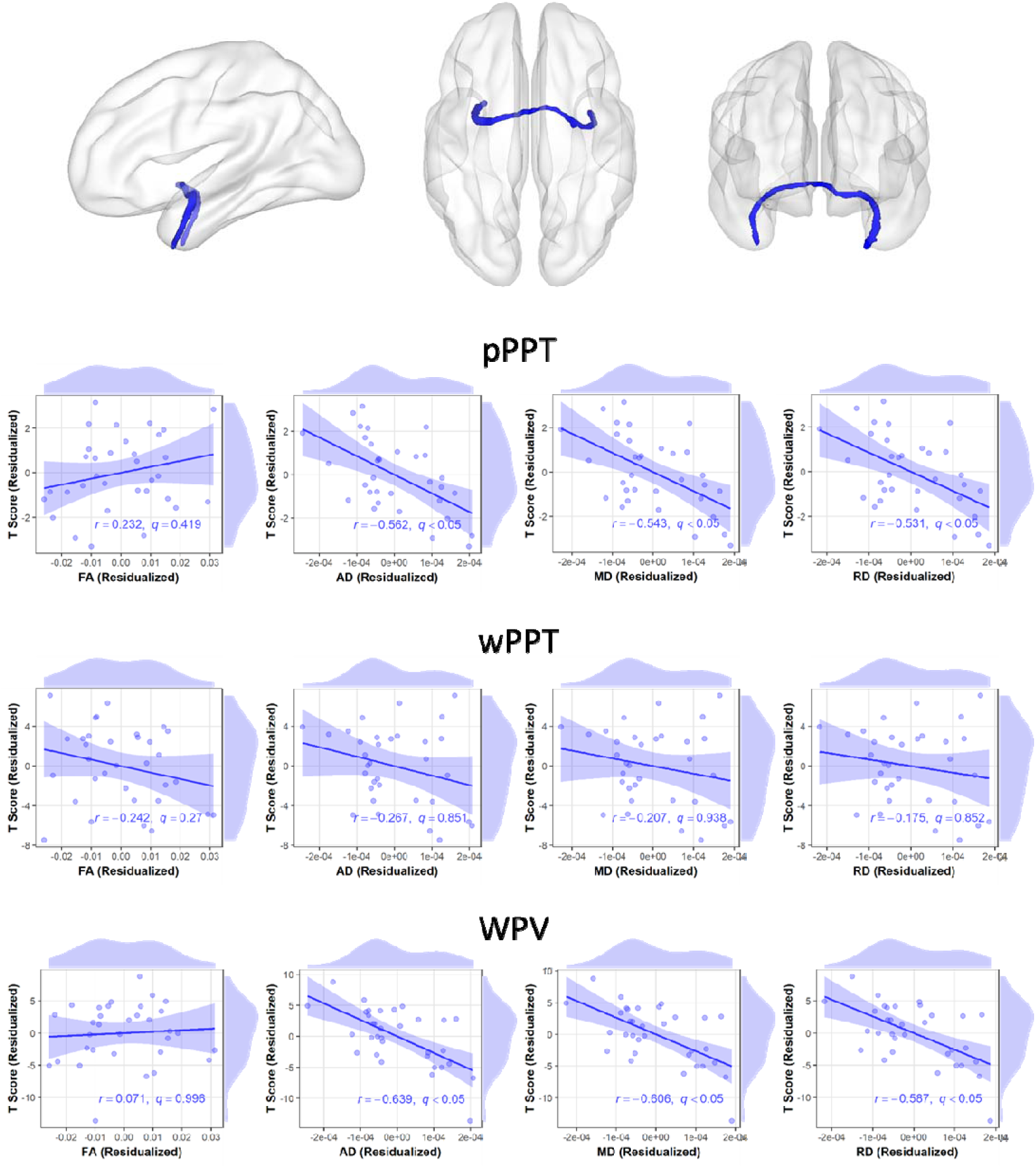
Correlation results between the integrity of the right ATL–left FFG white matter tract and performance on semantic processing tasks (WPV, pPPT, and wPPT) within the semantic dementia patient group, after regressing out TIV, performance on non-semantic control tasks, and lower-level language processing tasks. Raw *p* values are reported.

The white matter connection between the right ATL and the left ATL can also significantly predicted WPV performance for AD (*r* = −0.478, FDR-corrected *q* < 0.05). However, for MD and RD, the associations were only marginal (MD: *r* = −0.441, FDR-corrected *q* = 0.066; RD: *r* = −0.418, FDR-corrected *q* = 0.090), see Figure 8. Moreover, when additionally regressing out the GMV of the left FFG, this white matter connection remained a significant predictor of WPV performance for AD (*r* = −0.492, *p* < 0.05). We also noted that the white matter tract connecting the right ATL to the left FFG and the tract connecting the right ATL to the left ATL share similarities in the left temporal lobe, as both originate from the anterior portion of the left ATL or the anterior part of the left FFG. This suggests that the anterior temporal region serves as an important hub for white matter connections involved in semantic processing.

**Figure 8:**
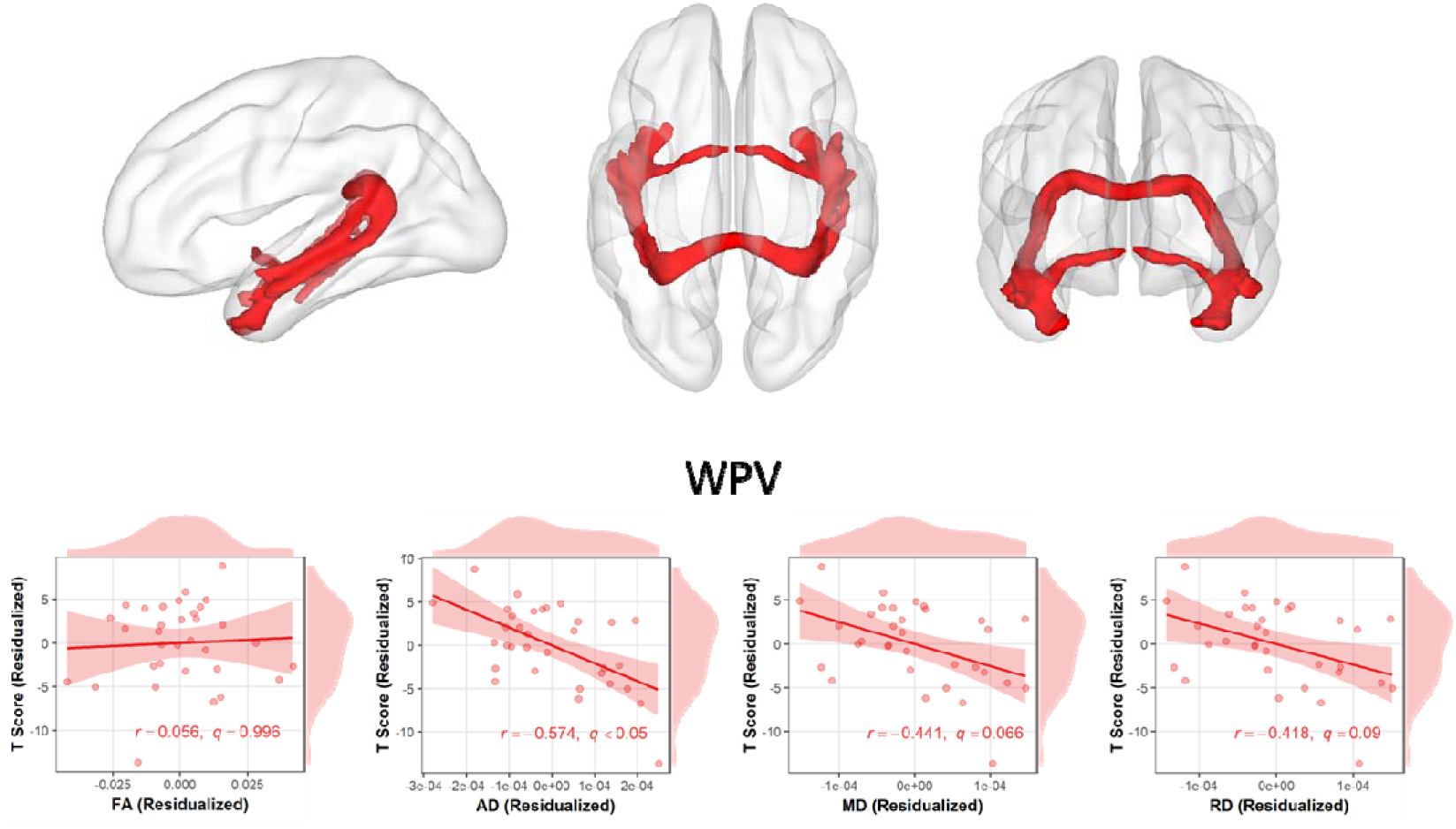
Patrial correlation results between the integrity of the right ATL–left ATL white matter tract and performance on WPV task within the semantic dementia patient group, after regressing out TIV, performance on non-semantic control tasks, and lower-level language processing tasks. Raw *p* values are reported.

None of the other white matter tracts showed significant predictive effects on performance across the three semantic processing tasks (FDR-corrected *q*s > 0.10), even after regressing out TIV, performance on non-semantic control tasks, and lower-level language processing tasks. Consequently, we did not report the additional partial correlation analyses that further controlled for GMV.

#### The role of white matter integrity in the integration of verbal and non-verbal semantic processing

For further analysis, we focused on the white matter tracts significantly associated with WPV performance in the previous step: the right ATL–left FFG and right ATL–left ATL connections. When TIV, as well as performance on the wPPT and pPPT tasks, were included as covariates, the strength of the white matter connection between the right ATL and the left FFG remained a significant predictor of WPV task performance (AD: *r* = −0.431, *p* < 0.05; MD: *r* = −0.424, *p* < 0.05; RD: *r* = −0.418, *p* < 0.05). Furthermore, this predictive effect remained significant even after additionally controlling for the GMV of the left FFG (AD: *r* = −0.384, *p* < 0.05; MD: *r* = −0.391, *p* < 0.05; RD: *r* = −0.393, *p* < 0.05). Moreover, the strength of the white matter connection between the right ATL and the left ATL was also a significant predictor of WPV task performance (AD: *r* = −0.476, *p* < 0.05; MD: *r* = −0.481, *p* < 0.05; RD: *r* = −0.479, *p* < 0.05). Importantly, this predictive effect remained significant even after additionally controlling for the GMV of the left FFG (AD: *r* = −0.512, *p* < 0.05; MD: *r* = −0.512, *p* < 0.05; RD: *r* = −0.507, *p* < 0.05), see Figure 9.

**Figure 9:**
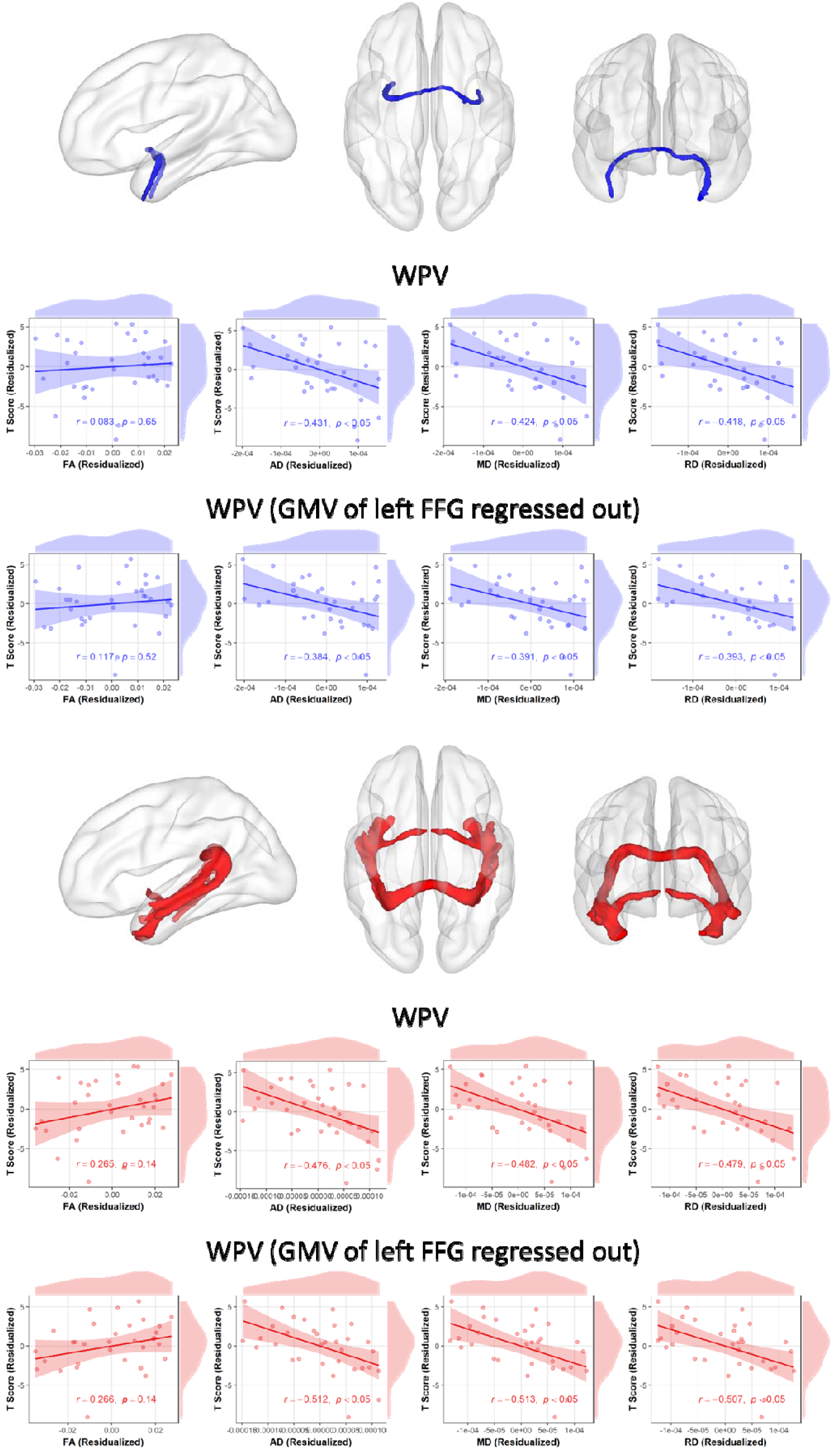
Partial correlation results between the integrity of the right ATL–left FFG (blue) and right ATL–left ATL (red) white matter tracts and WPV task performance within the semantic dementia patient group, after regressing out TIV, wPPT task performance, and pPPT task performance, as well as the additional results after regressing out the GMV of the left FFG. Raw *p* values are reported.

### Resting-state functional indices–behavior mapping analysis based on template-defined ROIs

#### ROI-based local functional indices–behavioral performance partial correlation analysis in the left hemisphere

Within the patient group, after regressing out TIV, performance on non-semantic control tasks (SP, VP, and NPM), and lower-level language processing tasks (reading and repetition), no local functional indices (ReHo, ALFF, fALFF, and DC) in the gray matter ROIs in the left hemisphere were found to significantly predict performance on the three semantic tasks, FDR-corrected *q*s > 0.10, see Table 3, 4 and 5.

**Table 3.** Partial Correlations Between Semantic Task Performance (wPPT) and Local Functional Indices in Patients.

| | Regressors | $r$ | FDR-corrected $q$ |
| --- | --- | --- | --- |
| **ReHo** |  |  |  |
| Left ATL |  | 0.127 | 0.787 |
| Left FFG |  | 0.055 | 0.787 |
| Left pMTG |  | 0.182 | 0.787 |
| **ALFF** |  |  |  |
| Left ATL | TIV, Performance | 0.082 | 0.762 |
| Left FFG | on SP, VP, NPM, | -0.061 | 0.762 |
| Left pMTG |  | -0.096 | 0.762 |
| **fALFF** |  |  |  |
|  | Reading and |  |  |
| Left ATL | Repetition | -0.178 | 0.510 |
| Left FFG |  | 0.260 | 0.510 |
| Left pMTG |  | 0.133 | 0.510 |
| **DC** |  |  |  |
| Left ATL |  | -0.022 | 0.915 |
| Left FFG |  | -0.144 | 0.712 |
| Left pMTG |  | 0.375 | 0.161 |

**Table 4.**
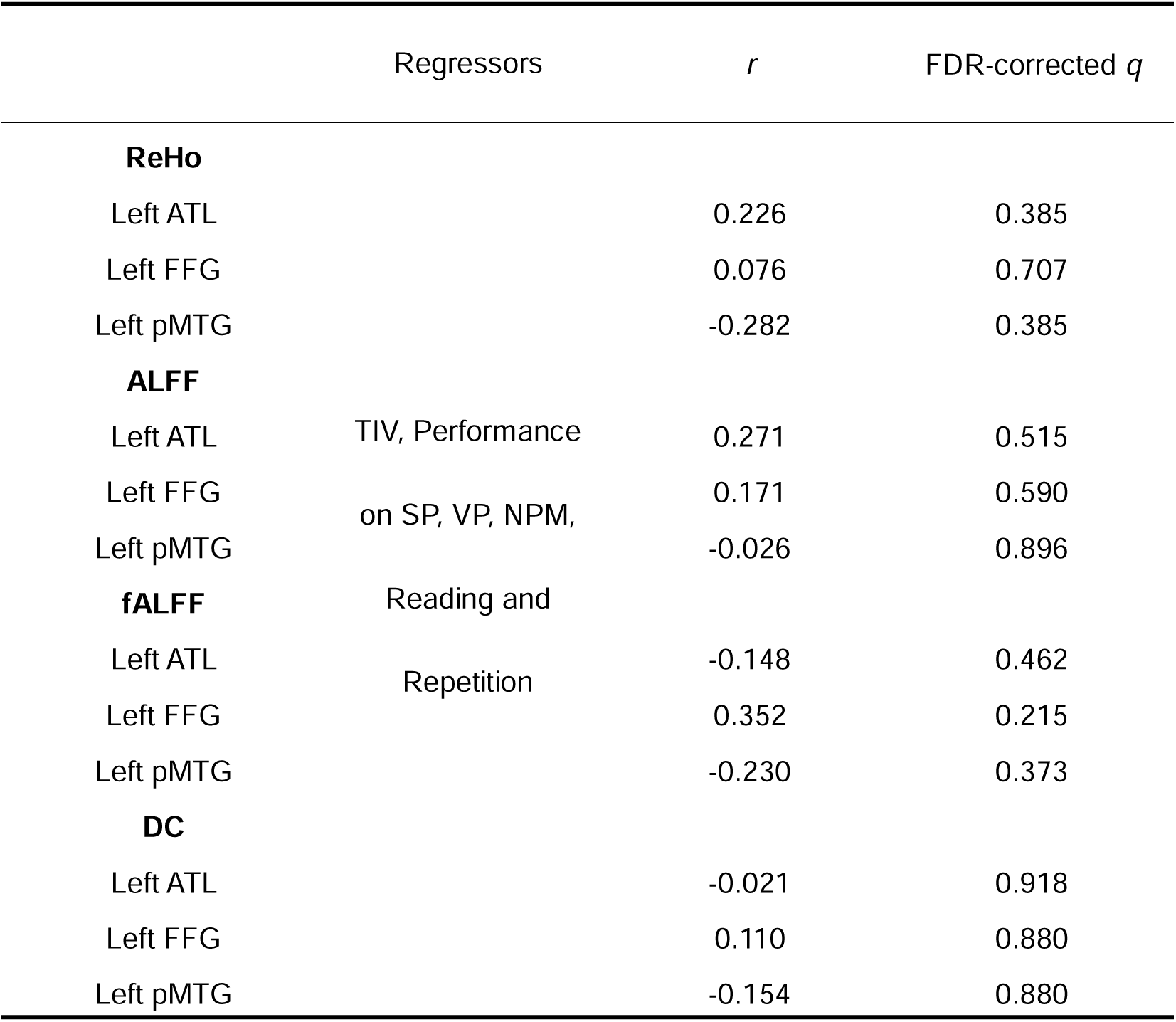
Partial Correlations Between Semantic Task Performance (pPPT) and Local Functional Indices in Patients.

**Table 5.**
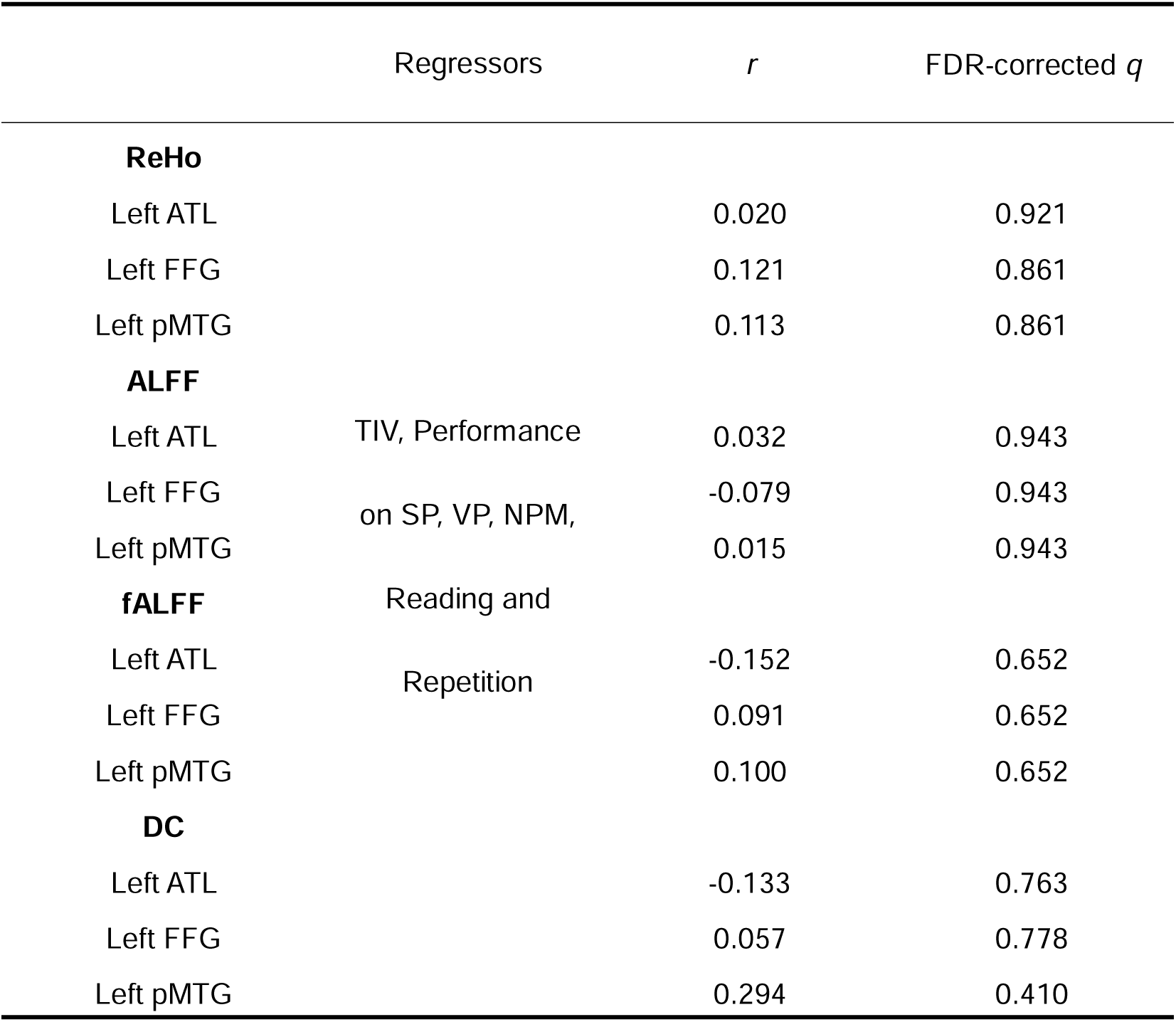
Partial Correlations Between Semantic Task Performance (WPV) and Local Functional Indices in Patients.

Since no significant associations were found between resting-state functional indices and performance on semantic processing tasks in the left hemisphere, we did not further examine the predictive effects of these indices on performance in the semantic integration task (WPV) after regressing out TIV and scores on the wPPT and pPPT tasks.

#### ROI-based local functional indices–behavioral performance partial correlation analysis in the right hemisphere

Within the patient group, after regressing out TIV, performance on non-semantic control tasks (SP, VP, and NPM), and lower-level language processing tasks (reading and repetition), no local functional indices (ReHo, ALFF, fALFF, and DC) in the right hemisphere gray matter ROIs were found to significantly predict performance on the wPPT or pPPT tasks (all FDR-corrected *q*s > 0.10). However, ALFF in the right FFG was found to significantly predict WPV task performance in a negative direction (*r* = −0.531, FDR-corrected *q*s < 0.05), whereas DC in the right pMTG significantly predicted WPV performance in a positive direction (*r* = 0.474, FDR-corrected *q*s < 0.05), see Table 6, 7 and 8.

**Table 6.**
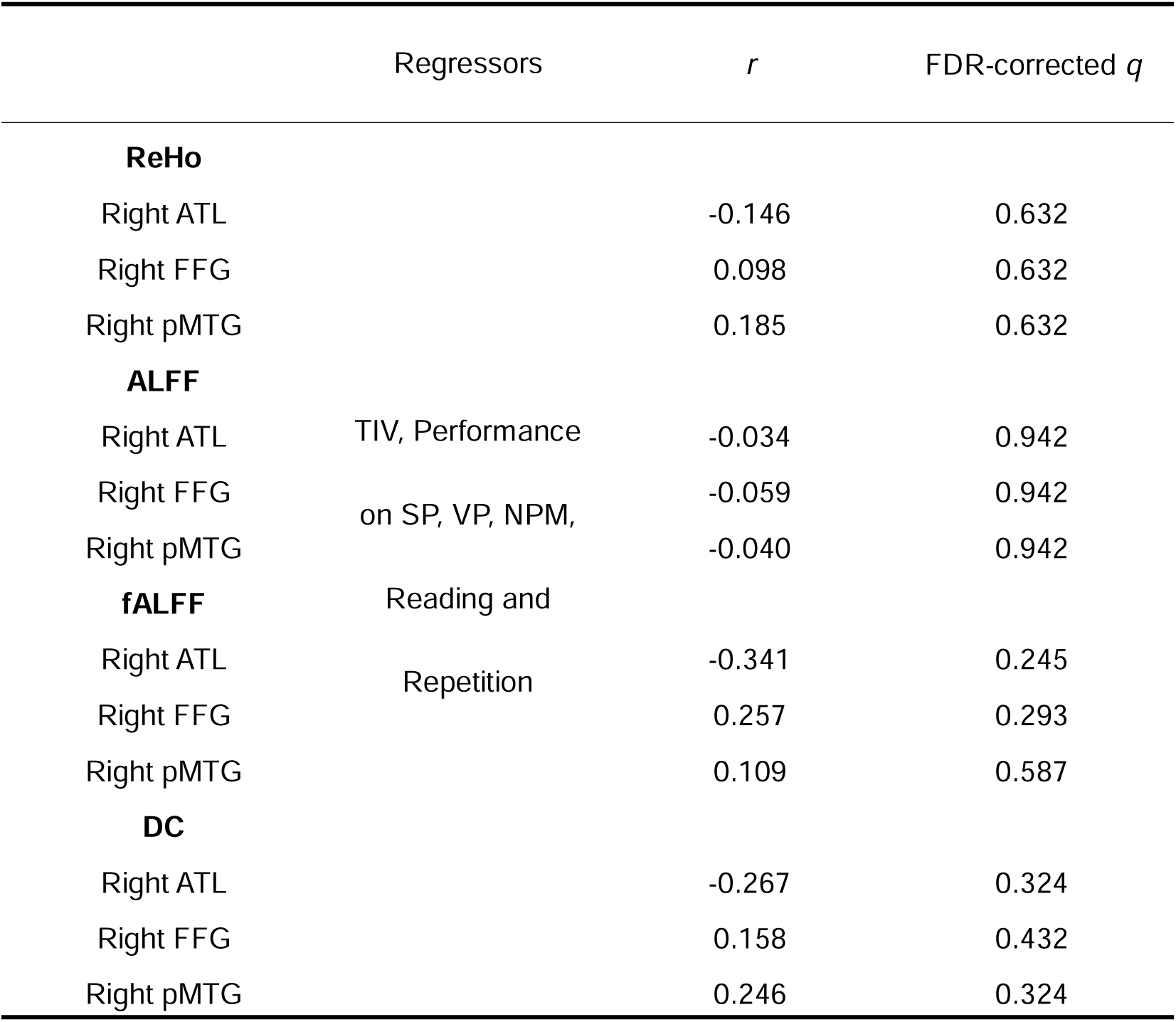
Partial Correlations Between Semantic Task Performance (wPPT) and Local Functional Indices in Patients.

**Table 7.** Partial Correlations Between Semantic Task Performance (pPPT) and Local Functional Indices in Patients.

Functional Indices in Patients
| | Regressors | $r$ | FDR-corrected $q$ |
| --- | --- | --- | --- |
| **ReHo** |  |  |  |
| Right ATL |  | 0.138 | 0.492 |
| Right FFG |  | 0.200 | 0.489 |
| Right pMTG |  | 0.355 | 0.208 |
| **ALFF** |  |  |  |
| Right ATL | TIV, Performance | < 0.001 | 0.999 |
| Right FFG | on SP, VP, NPM, | -0.410 | 0.101 |
| Right pMTG |  | -0.241 | 0.338 |
| **fALFF** | Reading and |  |  |
| Right ATL | Repetition | 0.024 | 0.904 |
| Right FFG |  | 0.213 | 0.431 |
| Right pMTG |  | 0.224 | 0.431 |
| **DC** |  |  |  |
| Right ATL |  | -0.050 | 0.805 |
| Right FFG |  | -0.229 | 0.749 |
| Right pMTG |  | 0.088 | 0.805 |

**Table 8.**
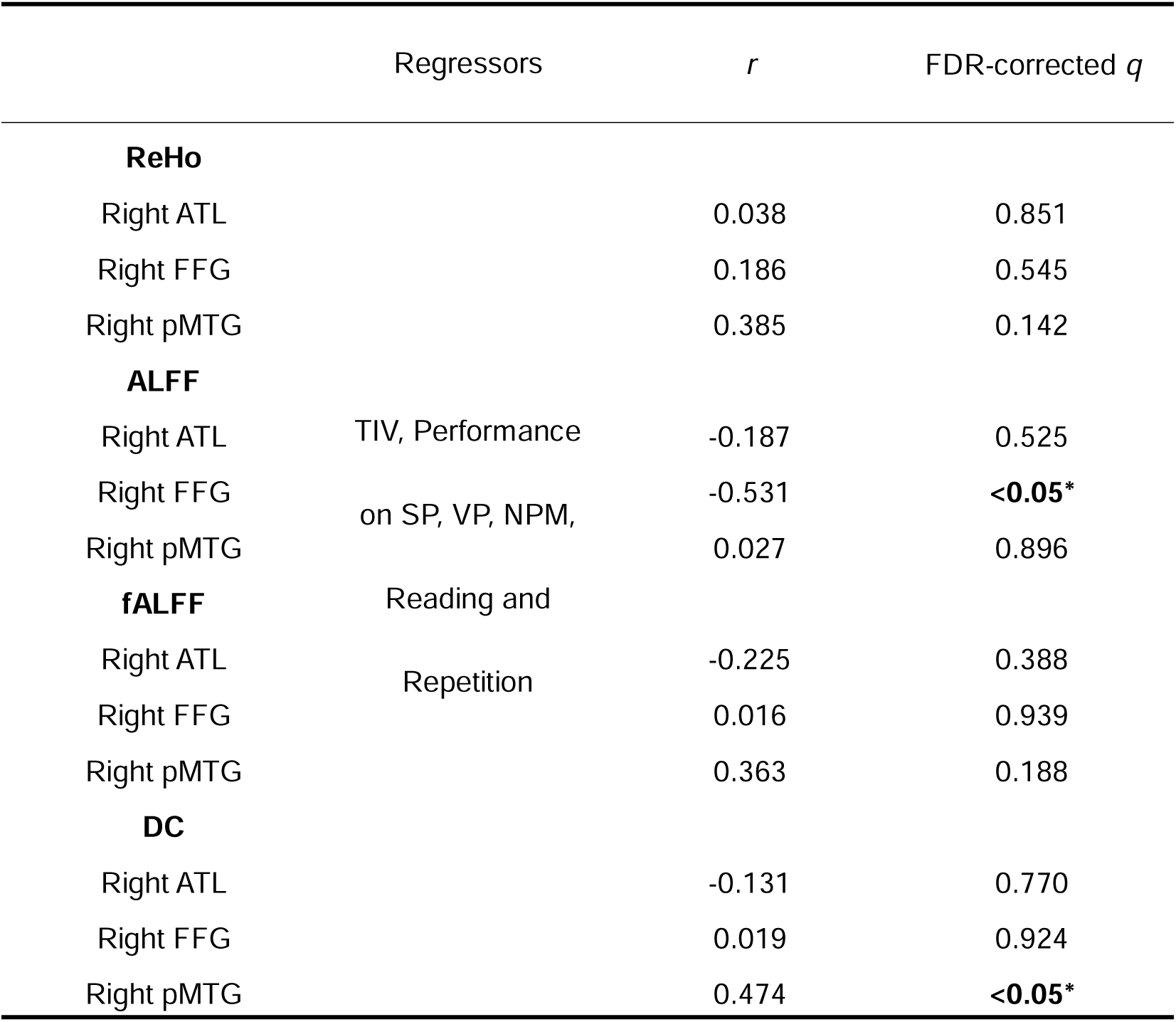
Partial Correlations Between Semantic Task Performance (WPV) and Local Functional Indices in Patients.

Additionally, when TIV, performance on wPPT, and pPPT tasks were included as covariates, the negative predictive effect of ALFF in the right FFG on WPV performance remained significant (*r* = −0.581, *p* < 0.05). In contrast, the positive predictive effect of DC in the right pMTG on WPV performance was only marginally significant (*r* = 0.338, *p* = 0.068).

## Discussion

We employed multimodal neuroimaging data to investigate the specific roles of six regions—three in each hemisphere (ATL, FFG, and pMTG)—commonly considered key hubs for semantic processing, encompassing both verbal and non-verbal domains and information integration. Analyses were conducted using partial correlation methods on data from individuals with semantic dementia. The findings reveal that the structural integrity of the left ATL is critical for verbal semantic processing; the left FFG contributes to both verbal and non-verbal semantic processing, suggesting its role as a core hub for general semantic processing; and the left pMTG appears to facilitate the integration of verbal and non-verbal semantic information. Although the structural integrity of the right ATL was not directly associated with semantic performance, its white matter connections with the left ATL and FFG were essential for semantic integration, highlighting a cross-hemispheric compensatory mechanism. This compensatory effect was further reflected in functional measures, including local neural activity and functional connectivity within the right FFG and pMTG. Moreover, these findings demonstrate varying sensitivity across imaging modalities, underscoring the importance of multimodal approaches for capturing region-specific contributions in semantic neuroscience.

### Left ATL and verbal semantic processing

This investigation found that the left ATL primarily supports verbal semantic processing, suggesting a specific functional specialization for abstract information processing. Within the classical “hub-and-spoke” framework, this region is considered a semantic hub that integrates inputs from multiple modalities, including visual, auditory, and tactile channels (Binney et al., 2016; Muraki et al., 2025; Patterson et al., 2007; Lambon Ralph et al., 2017). Damage to the ATL is known to cause severe impairments in abstract concept comprehension (Binney & Lambon Ralph, 2015; Chiou & Lambon Ralph, 2019; Jackson et al., 2015; Jefferies et al., 2009; Loiselle et al., 2012; Mollo et al., 2017b; Pobric et al., 2009). Our findings not only reinforce the view of the left ATL as a semantic hub but further indicate that its role may be particularly critical for extracting abstract verbal semantic representations from external stimuli (Jung & Lambon Ralph, 2021; Sierpowska et al., 2019). In other words, the left ATL may constitute a core neural substrate enabling humans to use language as a cognitive tool for understanding the world. The functional organization of this region is likely shaped by individual language experience and cognitive styles that emphasize linguistic thought (Bi et al., 2011; Wang et al., 2023). Compared to other mammals, humans exhibit a pronounced expansion of this area—a structural adaptation that underpins our unique ability to process abstract semantic information through language (Bi, 2021; Jung et al., 2025; Vignali et al., 2023; Wang et al., 2020). This suggests that the ATL serves as a critical structural foundation for abstract semantic cognition.

### Left pMTG and verbal and non-verbal semantic integration

The findings of this study further indicate that the left pMTG plays a critical role in semantic integration between verbal and non-verbal visual stimuli. Successful integration requires individuals to possess intact verbal and non-verbal semantic representations, suggesting that this region is involved in a process that goes beyond a single verbal or non-verbal semantic representation(Davey et al., 2016; Hoffman et al., 2012; Zhao et al., 2021) and likely occurs at a later stage in the cognitive sequence (Eisenhauer et al., 2024; Visser et al., 2012). In contemporary cognitive neuroscience, the pMTG is often considered part of the extended Wernicke’s area and has been identified in stroke research as an important region for language comprehension (Binder, 2017; Naeser et al., 1987; Tremblay & Dick, 2016; J. Wang et al., 2015). However, recent studies on patients with semantic dementia or primary progressive aphasia (PPA) have suggested that Wernicke’s area plays only a supportive role for ATL function in language comprehension (Matchin et al., 2022, 2023; Mesulam et al., 2015, 2019, 2023; Ogar et al., 2011). Although these studies have proposed that the pMTG may contribute to relatively low-level aspects of language processing, while the left ATL is implicated in higher-order and core components of word semantic comprehension, they have not clearly defined its specific supportive function. The present findings offer an alternative interpretation: the left pMTG may contribute to a later, higher-level process of cross-modal semantic integration, which builds upon the representational functions of the left ATL and therefore depends on the integrity of ATL function. This perspective may help reconcile the apparent discrepancy between findings from stroke research and studies of semantic dementia and PPA. Although Wernicke’s aphasia has traditionally been attributed to damage in the posterior superior temporal gyrus (pSTG), post-stroke lesions are often extensive and frequently involve the pMTG (Fridriksson et al., 2018; Knepper et al., 1989; Matchin et al., 2022; Turkstra, 2018). Damage to the pMTG may directly disrupt the integration of verbal and non-verbal semantic information, thereby contributing to impairments in language comprehension and semantic processing. In contrast, in semantic dementia, severe degradation of verbal semantic representations due to left ATL atrophy may further lead to diaschisis effects on the pMTG’s ability to integrate verbal and non-verbal information. From this perspective, the findings of the present investigation may provide a potential explanatory framework for the so-called “Wernicke Conundrum.” Therefore, even without relying on the potentially ambiguous and historically variable definition of Wernicke’s area, the results of this study suggest that posterior regions of the temporal lobe—specifically the left pMTG—should be considered core and high-level regions for language comprehension and semantic processing.

### Left FFG and general semantic processing

Extensive prior research has emphasized the fusiform gyrus (FFG) as a key region involved in general semantic processing (Ding et al., 2016; Forseth et al., 2018; Spagna et al., 2020; Zhang et al., 2016). Data-driven analyses based on diffusion tensor imaging (DTI) further support the role of the FFG as a semantic hub (Chen et al., 2020). In this study, we examined the relationship between structural damage to the FFG and performance on both verbal and non-verbal semantic tasks. Our findings demonstrate that the FFG contributes significantly to semantic processing across verbal and non-verbal stimuli, reinforcing its role in general semantic cognition. Although damage to the FFG predicted performance on semantic integration tasks involving both verbal and non-verbal stimuli, this effect was largely accounted for by atrophy in the left pMTG. A subsequent commonality analysis confirmed that the pMTG, rather than the FFG, primarily explained the variance in performance on the verbal–non-verbal semantic integration task (i.e., the WPV task), underscoring the critical role of the pMTG in cross-format semantic integration. These findings suggest that while the left FFG is involved in processing semantic information from both verbal and non-verbal visual stimuli, the integration of these two types of information relies predominantly on the left pMTG, highlighting the unique contribution of pMTG gray matter integrity to verbal and non-verbal semantic integration.

### Structural–functional interactions underlying verbal and non-verbal semantic integration

Partial correlation analyses of white matter integrity and behavioral performance revealed that the strength of white matter connections between the right ATL and the left ATL and FFG was significantly associated with verbal and non-verbal semantic integration processes. As shown in Figure 2, the fiber tract connecting the right ATL and the left FFG originates from the anterior portion of the FFG, highlighting the importance of anterior temporal regions within the white matter network. These findings suggest that although the ATL does not directly influence verbal and non-verbal semantic integration through gray matter integrity, it can exert an effect via the structural integrity of white matter connections, thereby impacting higher-order cognitive processes and revealing the complexity of interactions within the brain’s structural network. Combined with resting-state data, local neural activity in the right FFG was significantly negatively correlated with WPV task performance, while functional connectivity of the right pMTG showed a marginally significant positive correlation. This pattern may reflect structural miswiring: gray matter damage in the left temporal lobe disrupts the white matter connections between the left and right ATL, leading to impaired interhemispheric communication and altered neural activity in the right temporal lobe (Chen et al., 2018; Kleinhans et al., 2025; Mesulam, 2012, 2023). The right FFG, affected by such miswiring, exhibits compensatory hyperactivation or network imbalance, which fails to effectively support semantic integration and may even interfere with processing (Catani & Mesulam, 2008; Ding et al., 2016; Xiao et al., 2024). In contrast, the pMTG, located in the posterior temporal lobe, is less affected by miswiring and can provide partial compensation for semantic integration through functional networks, although insufficient to restore normal performance (Davey et al., 2015, 2016; Kocsis et al., 2023). These reveal the functional interdependence and dynamic influence between bilateral semantic hubs in the brain.

### Limitations

This study has several limitations that should be acknowledged: (1) It lacks data from other variants of PPA. In addition to semantic dementia, which typically involves atrophy originating in the ATL, this study did not include other PPA subtypes with different atrophy patterns, such as logopenic variant PPA, which often begins with atrophy in the posterior temporal regions. (2) This investigation focused exclusively on verbal and non-verbal semantic processing under visual input conditions. Future research should compare non-verbal stimuli presented visually with verbal stimuli presented auditorily, which would be crucial for testing whether the pMTG supports cross-modal integration of verbal and non-verbal semantic processing. (3) The absence of longitudinal data limits causal inference. Tracking patients over time with repeated assessments of semantic task performance and structural imaging would provide stronger evidence for establishing causal relationships.

## Conclusion

In conclusion, the findings of this investigation support the view that distinct semantic hubs exhibit functional specialization depending on the type of semantic information being processed. Specifically, the left ATL is primarily engaged in abstract verbal semantic processing; the left FFG contributes to general semantic processing across both verbal and non-verbal stimuli; and the left pMTG plays a central role in integrating verbal and non-verbal semantic information. Moreover, these hubs may influence contralateral regions through white matter connections, potentially driving changes in neural activity and functional network reorganization. Taken together, these findings underscore that even within the same input modality, different types of semantic stimuli recruit distinct semantic hubs, highlighting the functional specialization of the semantic network.

## Data availability statement

All data and study materials are available from the corresponding author on reasonable request.

